# Differential transcriptional activity of ΔNp63β is encoded by an isoform-specific C-terminus

**DOI:** 10.1101/2024.12.03.626646

**Authors:** Abby A. McCann, Morgan A. Sammons

## Abstract

p63 is a clinically-relevant transcription factor heavily involved in development and disease. Mutations in the p63 DNA-binding domain lead to severe developmental defects and overexpression of p63 plays a role in the progression of epithelial-associated cancers. Unraveling the specific biochemical mechanisms underlying these phenotypes is made challenging by the presence of multiple p63 isoforms and their shared and unique contributions to development and disease. Here, we explore the function of the p63 isoforms ΔNp63ɑ and ΔNp63β to determine the contribution of C-terminal splice variants on known and unique molecular and biochemical activities. Using RNA-seq and ChIP-seq on isoform-specific cell lines, we show that ΔNp63β regulates both canonical ΔNp63ɑ targets and a unique set of genes with varying biological functions. We demonstrate that the majority of genomic binding sites are shared, however the enhancer-associated histone modification H3K27ac is highly enriched at ΔNp63β binding sites relative to ΔNp63ɑ. An array of ΔNp63β C-terminal mutants demonstrates the importance of isoform-specific C-terminal domains in regulating these unique activities. Our results provide novel insight into differential activities of p63 C-terminal isoforms and suggest future directions for dissecting the functional relevance of these and other transcription factor isoforms in development and disease.

## INTRODUCTION

The transcription factor p63, encoded by the *TP63* gene, is essential for the development and homeostasis of the epidermis and epithelial-derived tissues (Fisher et al., 2020; Di Girolamo et al., 2024). *TP63* knockout mice exhibit severe craniofacial, limb, and epidermal defects, leading to neonatal lethality (Yang et al., 1998, p. 63; Mills et al., 1999). Heterozygous mutations in the DNA binding domain of *TP63* are linked to several human disorders including Ectrodactyly, Ectodermal dysplasia and Cleft lip/palate (EEC) (Celli et al., 1999; van Bokhoven et al., 2001). Mutations across other *TP63* exons result in a range of disorders with underlying dysfunction in epithelial cell biology (Di Girolamo et al., 2024). Consistent with the observed organism-level phenotypes, p63 transcription factor activity is required for epithelial lineage commitment and self-renewal (Li et al., 2023). These activities include interaction with gene regulatory elements like enhancers and promoters, control of local and long-distance chromatin structure, and transcriptional regulation of a pro-epithelial gene expression network (Fessing et al., 2011; Kouwenhoven et al., 2015; Li et al., 2019; Lin-Shiao et al., 2019). EEC patient-derived keratinocytes display dysregulated epidermal and epithelial-specific genes and an altered regulatory element landscape (Qu et al., 2018). Thus, understanding the mechanisms of gene regulation and molecular activities of p63 is crucial due to its significant impact on epithelial biology and human health.

*TP63* is expressed as several isoforms, through a combination of alternative promoter usage and alternative C-terminal splicing. The major isoforms include two N-terminal variants, TA and ΔN, and at least four C-terminal splice variants (ɑ,β,γ,Δ), yielding 8 isoforms (Murray-Zmijewski et al., 2006; Sethi et al., 2015; Marshall et al., 2021). An additional N-terminal isoform, GTAp63ɑ, contains an elongated N-terminal domain relative to TAp63 and is predominantly expressed in male germ cells (Pitzius et al., 2019). Most prior analyses of p63 function primarily focused on the TAp63ɑ and ΔNp63ɑ isoforms. TAp63ɑ, expressed in oocytes and during late keratinocyte differentiation, performs p53-like functions in maintaining genome integrity by inducing apoptosis after DNA damage (Suh et al., 2006; Livera et al., 2008). Studies using knockout mice and *in vitro* cell culture approaches have shown that ΔNp63ɑ is the N-terminal isoform primarily responsible for developmental and epithelial-related defects and for controlling epithelial-related gene and chromatin networks (Mills et al., 1999; Yang et al., 1999; Suh et al., 2006; Su et al., 2009; Wolff et al., 2009). Further highlighting the importance of p63ɑ isoforms, the human disorder Ankyloblepharon-ectodermal defects-cleft lip/palate (AEC) is caused by heterozygous mutations in the alpha-specific SAM domain (McGrath et al., 2001). However, the specific contribution of individual ΔNp63 C-terminal isoforms to various developmental and transcriptional phenotypes is not fully resolved. Since most *TP63* knockout mouse models target the DNA binding domain, shared across all known isoforms, analysis of isoform-specific epithelial phenotypes or molecular activities has been complicated.

*In vitro* and cell-based analyses have identified some unique activities of the C-terminal p63 isoforms. The alpha-specific C-terminus inhibits transcription of the TA isoform and likely controls gene repression activities of ΔNp63ɑ (LeBoeuf et al., 2010; Ramsey et al., 2011; Coutandin et al., 2016; Pitzius et al., 2019). All other C-terminal variants lack this domain, including ΔNp63β, which has increased transcriptional activity *in vitro* and has increased anti-proliferative activity in cell models relative to other C-terminal isoforms (Helton et al., 2006, 2008). Meta-analysis of RNA-seq and other targeted expression analyses suggest that ΔNp63β is expressed in similar cell types as ΔNp63ɑ, albeit at lower levels (Sethi et al., 2015; Marshall et al., 2021). *In vivo,* ΔNp63β complements some ΔNp63ɑ function in *Trp63^-/-^* mice, such as transcriptional control of basal epithelial genes keratin 5 (K5) and keratin 14 (K14) (Romano et al., 2009). Mice heterozygous for deletion of *Trp63* exon 13, containing the SAM domain, have increased expression of ΔNp63β. Although there was no disruption of epidermal-related development, co-expression of ΔNp63ɑ and ΔNp63β led to female germ cell apoptosis and ovarian insufficiency presumably through increased transcriptional activity of ΔNp63β (Lena et al., 2021). Therefore, the limited experimental investigation into p63 C-terminal isoforms suggests both shared and isoform-specific unique activities, and the specific mechanisms that confer those differences are not fully explored.

We sought to further investigate p63 C-terminal variant activity, focusing on functional differences between ΔNp63ɑ and ΔNp63β. Our data suggest that ΔNp63β is capable of carrying out many canonical ΔNp63ɑ functions, but has increased transcriptional activity and a unique gene regulatory network. These differences in gene regulation are unlikely due to differences in genomic binding, but rather likely reflect differential activity at regulatory elements, including more widespread induction of enhancer-associated H3K27ac. Our data also provide evidence that ΔNp63β activity requires a protein domain shared with ΔNp63ɑ and ΔNp63Δ, but that is uniquely critical in ΔNp63β in combination with a β-specific 5 amino acid C-terminal domain. Thus, our data provide additional support for the observation that ΔNp63ɑ and ΔNp63β share limited roles in control of epithelial-related gene regulation and provide novel insight into the genomic and molecular mechanisms by which ΔNp63β may control unique biological functions through increased transcriptional activity.

## RESULTS

### The ΔNp63β isoform exhibits high transcriptional activity

N-terminal p63 isoforms encode two different N-terminal transactivation domains (TADs). The TAp63 isoforms have a well-characterized and highly active N-terminal TAD similar in structure to the p53 N-terminal TAD (Fig. 1A). ΔNp63 isoforms contain a unique 14 amino acid N-terminal region generated by an alternative transcriptional start site (Yang et al., 1998) (Fig. 1B). The absence of the canonical N-terminal TAD in ΔNp63 isoforms is thought to reduce transactivation relative to TA isoforms (Dohn et al., 2001), although the specific contribution of isoform-specific C-terminal domains to transcriptional control and observed biological activity is not fully characterized. To better understand the differences in function between C-terminal isoforms (Fig. 1A,B), we measured relative transcriptional activity of each p63 isoform using a reporter assay encoding a defined, synthetic p63 response regulatory element (RE). Each isoform, along with a negative control vector, was transfected into HCT116 *TP53^-/-^*colon carcinoma cells to avoid potential crosstalk with p53-dependent transcriptional activity (Fig. 1C,D). The sequence and GenBank/UniProt accession number for each isoform is available in Table S1. TA isoforms were all capable of activating transcription driven by the wild-type p63RE, but not a reporter containing a mutant p63RE (Fig. 1E). TAp63ɑ activated transcription over background levels, although its activity was at least 10-fold lower than the other three isoforms (Fig.1E). Activity of the TA β, γ, and Δ isoforms was similar, suggesting the absence of the TID is more important to their activity than the inclusion of any isoform-specific domains. These results are broadly consistent with prior work noting high activity of TAp63 isoforms and auto-inhibition of TAp63ɑ by the C-terminal inhibitory domain (TID). ΔNp63ɑ, ΔNp63γ, and ΔNp63Δ all exhibited similar levels of transactivation in contrast to the behavior of these C-terminal isoforms of TAp63 (Fig. 1F). ΔNp63β, however, was nearly 30-fold more active compared to the other ΔN isoforms. This β-specific increase in transactivation relative to ΔNp63γ, and ΔNp63Δ isoforms was not observed for TAp63β, suggesting a potentially unique mechanism driving activity of the ΔNp63β isoform.

**Figure 1:**
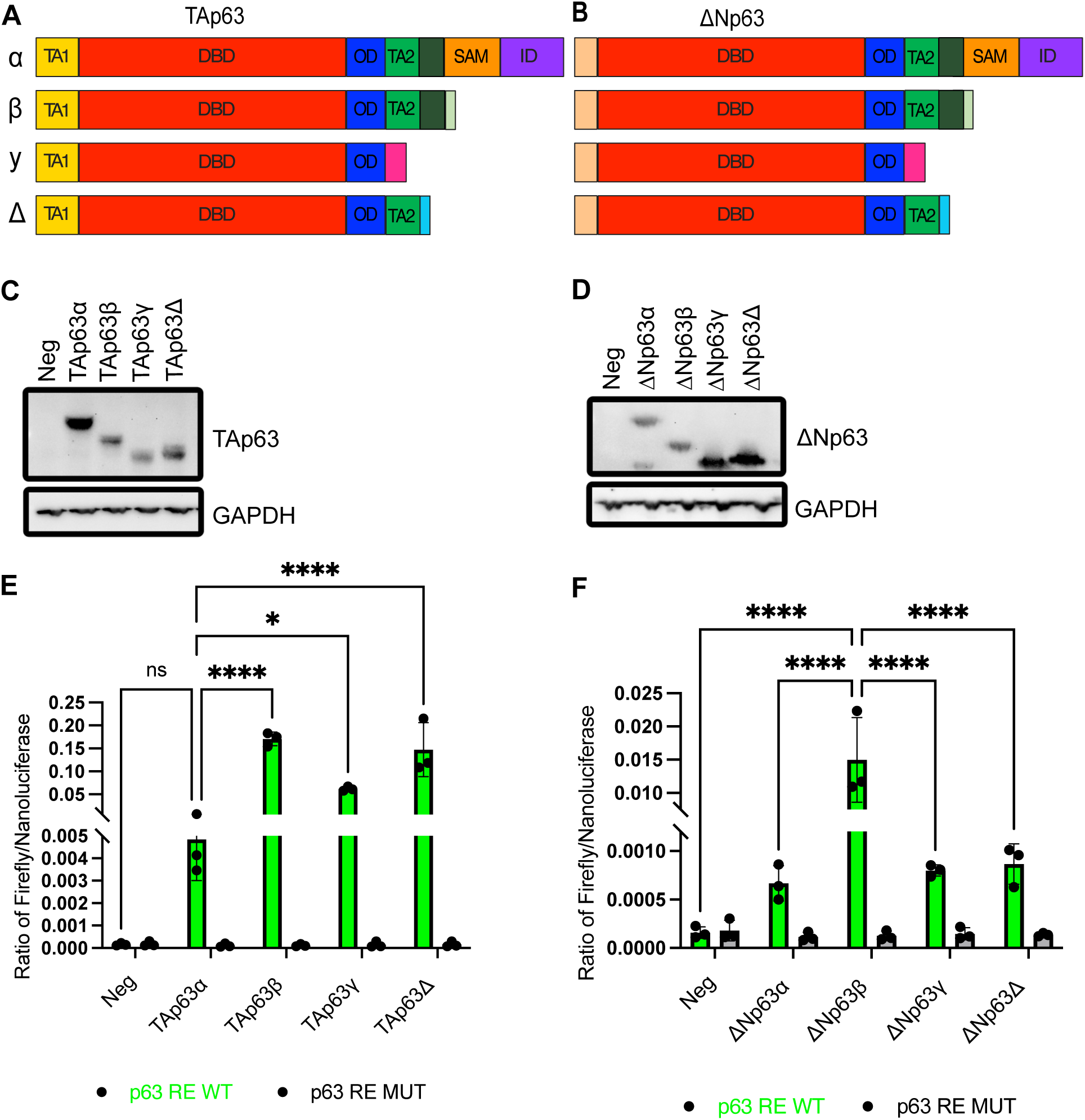
Relative transcriptional activity of the p63 isoforms. **A)** Schematic of the four TAp63 isoforms and the four **B)** ΔNp63 isoforms. **C)** Western blot protein expression of the TAp63 and **D)** ΔNp63 isoform constructs in pcDNA backbone transiently transfected into HCT116 *TP53^-/-^* cells using isoform specific antibodies. **E)** Reporter assay of the four TAp63 isoforms and the **F)** four ΔNp63 isoforms on a p63 responsive regulatory element (green) and a mutant (grey) p63 regulatory element in HCT116 *TP53^-/-^* cells. Negative control for western blot and reporter assay is an empty pcDNA backbone. (*: *p*-value <.05, ****: *p*-value < 0.0001, ns=not significant, Two-way ANOVA).

### RNA-seq analysis reveals shared and unique roles for ΔNp63β

Limited data are available comparing global gene expression programs controlled by p63 isoforms. The unique temporal and spatial expression patterns of these isoforms provide a challenge for determining their biological function and transcriptional regulation *in vivo* (Marshall et al., 2021). ΔNp63β has been reported to phenocopy certain roles of ΔNp63ɑ, but also has striking differences in transactivation potential (Fig. 1F). Therefore, we sought a better understanding of the differences between these two p63 isoforms by examining their gene regulatory potential. The likelihood that p63 isoforms can form mixed heterotetramers can complicate dissection of isoform-specific roles (Yin et al., 2002; Guo et al., 2024). Therefore, we performed bulk transcriptome profiling using RNA-seq to compare the differential gene expression between HCT116 *TP53^-/-^* cells expressing either ΔNp63ɑ or ΔNp63β under doxycycline inducible conditions (Fig. 2A). These cells do not natively express any p63 isoforms and also lack other p53 family members, thus any transcriptome changes can be more easily attributed to the specific isoform being expressed. PolyA+ RNA was isolated and sequenced using standard short-read, Illumina sequencing approaches. Transcript abundance within the ENSEMBL v.104 reference was quantified using *kallisto* and differential gene expression values (no induction vs. 24 hour doxycycline induction) were determined using DESeq2. Differentially expressed genes were called with a Bonferonni-corrected P-value of less than 0.05. Principal Component Analysis (PCA) demonstrates that biological replicates cluster together and that ΔNp63ɑ and ΔNp63β are in distinct clusters (Fig. S1A).

**Figure 2:**
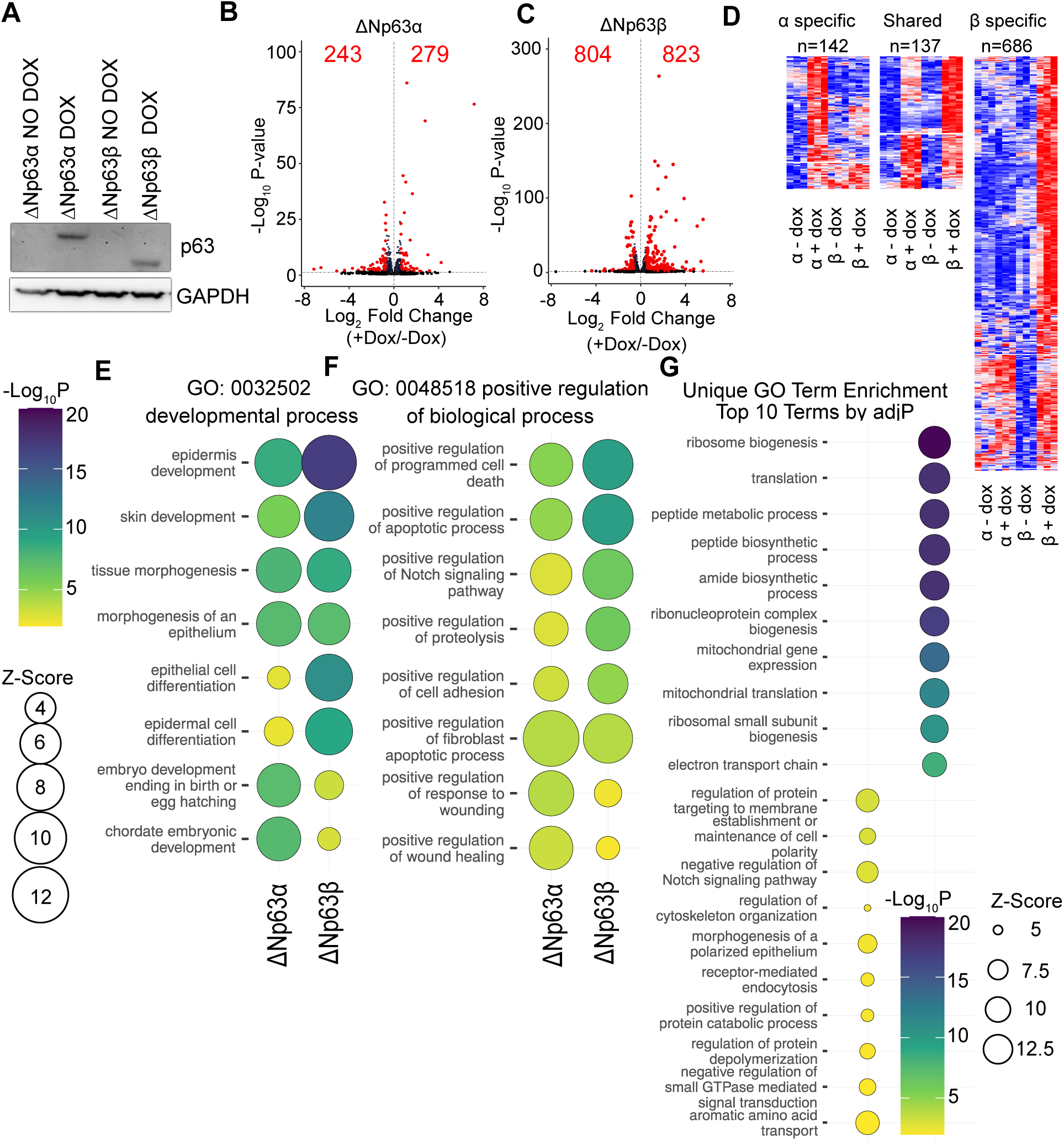
RNA-seq analysis identifies transcriptional targets of ΔNp63ɑ and ΔNp63β. **A)** Western blot for p63 (ΔN-specific domain) or GAPDH from HCT116 *TP53^-/-^* cells expressing either ΔNp63ɑ or ΔNp63β after 24 hour doxycycline induction. Volcano plot of differentially expressed genes after induction of **B)** ΔNp63ɑ or **C)**. Red points represent differentially expressed genes in induction conditions (+ doxycycline, 24hrs) relative to uninduced conditions (-doxycycline, 24hrs) at a Bonferroni-corrected P-value of less than or equal to 0.05. **D)** Heatmaps of differentially expressed genes (q-value <= 0.05) that are shared, or specific to either ΔNp63ɑ or ΔNp63β-induced conditions. **E)** Gene ontology enrichment of upregulated genes after induction of either ΔNp63ɑ or ΔNp63β, showing the top 10 child terms from the shared Developmental Process Parent Term. **F)** Gene ontology enrichment of upregulated genes after induction of either ΔNp63ɑ or ΔNp63β, showing the top 10 child terms from the shared Positive Regulation of Biological Processes Parent Term. **G)** Top 10 Gene Ontology terms uniquely identified in either ΔNp63ɑ or ΔNp63β differentially upregulated genes.

ΔNp63β induction leads to a greater number of differentially expressed genes relative to ΔNp63ɑ (Fig. 2B-C), consistent with increased ΔNp63β-dependent transcriptional activation in reporter assays (Fig. 1F). Gene ontology analysis of differentially upregulated genes shows shared functions in epidermis and skin development, tissue morphogenesis and epithelial and epidermal cell differentiation, suggesting ΔNp63β has the ability to carry out activities canonically associated with ΔNp63ɑ (Fig. 2E). A full list of gene ontology results can be found in Table S2. Both ΔNp63ɑ and ΔNp63β induce genes involved in other canonical p53 family activities, such as regulation of programmed cell death (Fig. 2F). ΔNp63ɑ and ΔNp63β downregulated genes involved in cell proliferation, cell migration, and epithelial cell differentiation (Fig. S1D), which supports a model whereby these two p63 isoforms share certain overlapping transcriptional roles.

Our data suggest that ΔNp63ɑ and ΔNp63β regulate a set of common gene targets (Fig. 2D, Fig. S1B) and that these genes are involved in canonical p63-dependent processes (Fig. 2E-F, Fig. S1D). The dramatic increase in ΔNp63β-specific genes (Fig. 2D, Fig. S1B) and a unique set of ΔNp63ɑ targets suggests these isoforms may also have key differences in target gene regulation and biological activity. 142 genes were uniquely upregulated by ΔNp63ɑ and 686 were unique to ΔNp63β (Fig. 2D). Unique GO terms for these differentially expressed genes for ΔNp63β show potential activities controlling anabolic processes, such as ribosome biogenesis, translation, and peptide synthesis (Fig. 2G). Uniquely downregulated ΔNp63β gene targets (Fig. S1B) were clustered into GO terms suggesting ΔNp63β-dependent control of cell cycle and cell division (Fig. S1C). GO terms associated with unique ΔNp63ɑ upregulated genes involve regulation of epithelial morphogenesis and cytoskeleton organization in addition to negative regulation of Notch signaling (Fig. 2G). p63 and Notch have a known, antagonistic relationship in epithelial cell regulation (Nguyen et al., 2006; Yalcin-Ozuysal et al., 2010; Tadeu and Horsley, 2013). Downregulated ΔNp63ɑ targets are associated with a range of unique Gene Ontology categories (Fig. S1C), including multiple groups suggesting control of epithelial cell and keratinocyte cell differentiation. Taken together, our analysis of ΔNp63ɑ and ΔNp63β-regulated genes suggests shared transcriptional roles in well-studied, p63-dependent processes but also unique transcriptional targets that may underlie isoform-specific biological activities.

### ChIP-seq analysis reveals predominantly shared binding sites for ΔNp63ɑ and ΔNp63β

Differential gene expression analysis suggests that ΔNp63ɑ and ΔNp63β regulate both an overlapping group of genes as well as isoform-specific targets. ΔNp63β regulates more genes than ΔNp63ɑ in our analysis. While consistent with increased transactivation by ΔNp63β in reporter assays (Fig. 1F), these observations may not fully explain differences in gene regulation. To further explore the mechanisms of the expanded ΔNp63β target gene network, we performed ChIP-seq of ΔNp63ɑ and ΔNp63β to ask whether differential regulation is linked to differential genomic binding (Fig. 3A). We performed these assays in HCT116 *TP53^-/-^*cells (Fig. 2A) to match the RNA-seq results and to prevent the possibility of endogenous ΔNp63ɑ confounding downstream analysis. Resulting ChIP-seq data were aligned to the hg38 reference genome using HiSat2 (Kim et al., 2019). Regions of enrichment were called relative to p63 ChIP from HCT116 *TP53^-/-^* expressing GUS, our negative control, using *macs2* and peaks from biological replicates were merged after filtering for locations within the hg38 blacklist regions (Amemiya et al., 2019). ΔNp63ɑ and ΔNp63β share 26,818 genomic binding sites (Fig. 3A). ΔNp63ɑ (10,185) and ΔNp63β (9.209) each have a unique set of binding sites, although these unique binding events are relatively low in enrichment in comparison to their shared binding sites (Fig. 3A-B). While ΔNp63ɑ is more enriched in binding sites called as unique in ΔNp63ɑ (Fig. 3A), ΔNp63β enrichment is present above the negative control background. Conversely, ΔNp63ɑ signal is enriched relative to negative background control in unique ΔNp63β binding sites. These data suggest ΔNp63ɑ and ΔNp63β bind to largely similar locations, but that isoform-specific preferences may drive higher enrichment at specific genomic loci.

**Figure 3:**
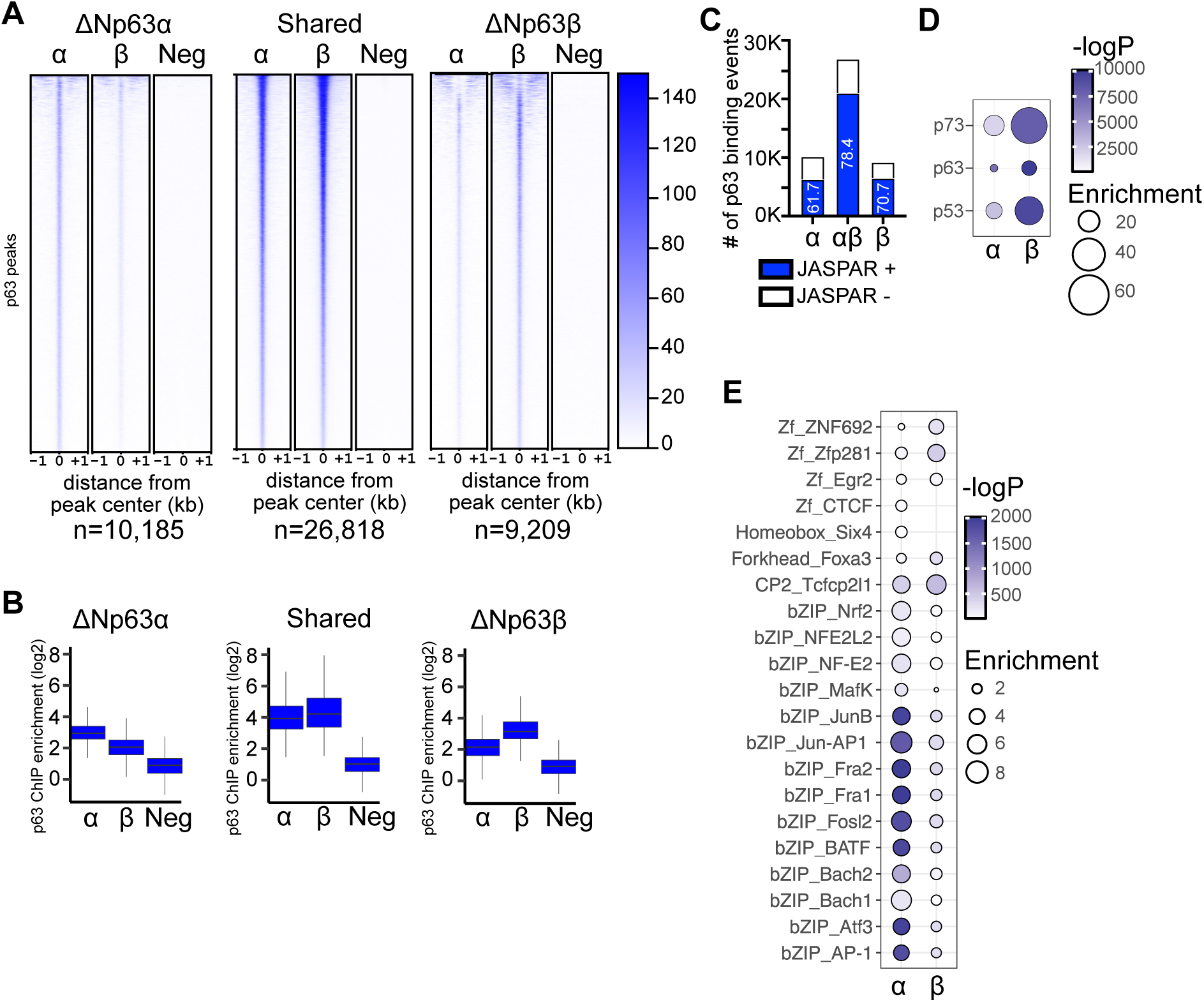
Genomic occupancy of ΔNp63ɑ and ΔNp63β. **A)** Heatmaps of ChIP-seq read density at MACS2-derived peaks found in only ΔNp63ɑ experiments, only ΔNp63β, or shared between both factors. Read densities (RPKM normalized) of ΔNp63ɑ, ΔNp63β, or empty vector negative control ChIP are plotted -/+ 1,000 base pairs from the peak center. Replicate data were merged into a single meta-plot. **B)** Quantification of read densities (log2 normalized) for ΔNp63ɑ, ΔNp63β, or IgG control ChIP-seq experiments at each class of peak. **C)** Percentage of shared or isoform-enriched ChIP-seq peaks containing a JASPAR-derived p53 family motif (p53, p63, or p73). **D**) HOMER-derived motif analysis for p53, p63, and p73 response elements from ΔNp63ɑ or ΔNp63β peak regions. Enrichment is relative to matched genomic background regions. **E)** HOMER-derived transcription factor motif enrichment from ΔNp63ɑ or ΔNp63β peak regions. Enrichment is relative to matched genomic background regions.

We next performed a series of DNA motif analyses to further characterize genomic binding preferences of ΔNp63ɑ and ΔNp63β. Canonical, JASPAR-derived p53 family motifs (p53,p63, and p73) are more common in shared binding sites compared to either ΔNp63ɑ or ΔNp63β-enriched locations (Fig. 3C). ΔNp63β binding sites are more highly enriched for p53, p63, and p73 motifs than ΔNp63ɑ using either JASPAR-defined motifs or when using HOMER to assess enrichment relative to genomic background (Fig. 3C-D). AP-1 family bZIP transcription factor motifs, commonly enriched in gene regulatory elements and associated with chromatin accessibility, are significantly more enriched in ΔNp63ɑ binding sites than in ΔNp63β (Fig. 3E). CTCF motifs are more commonly found in ΔNp63ɑ sites (Fig. 3E), consistent with prior reports of cooperation between ΔNp63ɑ and CTCF in gene regulation (Qu et al., 2019). Overall, our genomic occupancy and motif enrichment analyses demonstrate key differences in binding locations and transcription factor motif enrichment at ΔNp63ɑ and ΔNp63β binding sites. The differences in genomic occupancy and the presence of other TF motifs near p63 binding sites provides an additional potential mechanism underlying differential gene expression regulation by ΔNp63 C-terminal isoforms.

### The genomic occupancy of ΔNp63ɑ and ΔNp63β is associated with shared and unique regulation of gene expression

Although ΔNp63ɑ or ΔNp63β share considerable overlap in their genomic binding, we observe isoform-specific enrichment at a set of genomic locations. We asked if shared or isoform-specific ΔNp63ɑ and ΔNp63β binding locations were linked to specific groups of gene targets. We used the polyEnrich approach which links binding locations to genes and then performs gene set enrichment to determine whether these binding events cluster into related Gene Ontology categories (Welch et al., 2014; Lee et al., 2020). The Top 10 Gene Ontology Biological Process (GOBP) terms associated with shared binding sites of ΔNp63ɑ or ΔNp63β are primarily associated with development of the epithelium and morphogenesis (Fig. 4F), canonical activities of p63. These GO terms are also strongly enriched for ΔNp63ɑ and ΔNp63β-specific locations suggesting that these unique binding sites contribute to some well-known biological functions attributed to p63. The most statistically enriched gene sets associated with only shared sites contain some developmental and epithelial terms, but also multiple terms related to programmed cell death (Fig. 4G). Supporting the use of this approach linking binding events to gene sets, we observe highly similar gene ontology groups when examining genes induced by ΔNp63ɑ and ΔNp63β in our RNA-seq analysis (Fig. 2E-F). Gene sets uniquely linked to ΔNp63ɑ-specific binding are primarily associated with cell adhesion, protein transport, and cytoskeletal organization (Fig. 4H), biological processes also suggested to be regulated by ΔNp63ɑ in our RNA-seq analysis (Fig. 2F-G). polyEnrich analysis suggests ΔNp63β-specific binding is linked to genes related to cell cycle regulation and inflammatory/immune processes (Fig. 4I) (Lee et al., 2020). A full list of polyEnrich results is available in Table S3. Thus these data, combined with our prior RNA-seq-based gene set enrichment work, suggest that unique genomic locations for ΔNp63ɑ and ΔNp63β are associated with specific groups of genes with different biological activities. Further, shared ΔNp63ɑ and ΔNp63β binding events are linked to canonical p63-dependent activities like regulation of epithelial development and regulation of cell death and proliferation.

**Figure 4:**
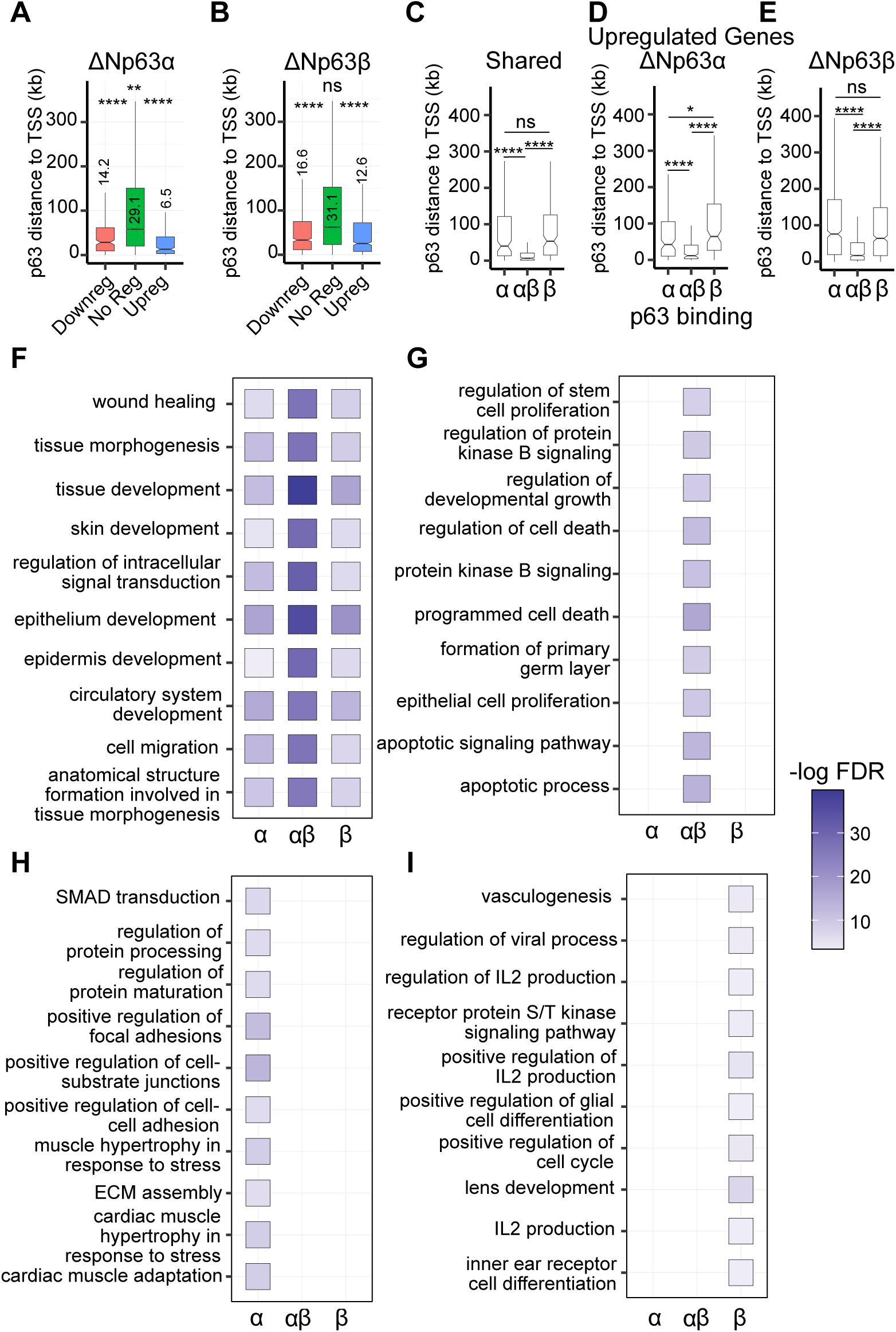
Integration of transcriptomes and cistromes reveals potential regulatory activities of ΔNp63ɑ and ΔNp63β. Analysis of the distance (in kilobases,kB) between either **A)** ΔNp63ɑ or **B)** ΔNp63β ChIP-seq peaks and the transcriptional start site of upregulated, downregulated, or unregulated genes after induced expression of each isoform. Regulated genes were classified as any fold-change relative to uninduced conditions with a Bonferonni-corrected P-value of less than 0.05 as determined by *DESeq2*. (**; p-value < 0.01, ****: *p*-value < 0.0001, Dunn’s Multiple Comparison Test). Analysis of the distance (in kilobases,kB) between shared, ΔNp63ɑ-enriched, or ΔNp63β-enriched ChIP-seq peaks and the transcriptional start site of **C)** shared, **D)** ΔNp63ɑ, or **E)** ΔNp63β-specific differentially regulated genes. (*; p-value < 0.05, ****: *p*-value < 0.0001, Dunn’s Multiple Comparison Test). Results of polyEnrich analysis of genes nearest shared, ΔNp63ɑ, or ΔNp63β ChIP-seq binding sites, displaying the top 10 (by FDR) Gene Ontology Biological Processes categories **F)** shared in all datasets, **G)** found only in shared binding events, or in either **H)** ΔNp63ɑ-specific sites **I)** ΔNp63β-specific sites. A full list of chipEnrich/polyEnrich results can be found in Table S3.

To further explore the link between binding of p63 isoforms and gene regulation, we asked whether the location and distance of either ΔNp63ɑ or ΔNp63β ChIP-seq binding sites to transcriptional start sites (TSS) of upregulated, downregulated or unchanged genes from the RNA-seq data (Fig. 3F) might correlate with differential gene regulation. For both ΔNp63ɑ (Fig. 4A) and ΔNp63β (Fig. 4B), ChIP-seq binding sites are significantly closer to TSS’s for both up and downregulated genes than unregulated genes. ΔNp63ɑ binding sites are closer to upregulated genes TSS than ΔNp63β (Fig. 4A vs. 4B), although whether this approximate 6kB difference is meaningful *in vivo* is unknown. We then asked whether the isoform-specific binding events are linked to specific transcriptional differences between ΔNp63ɑ and ΔNp63β. Genes activated by both ΔNp63ɑ and ΔNp63β are significantly closer to shared binding sites than either unique ΔNp63ɑ or ΔNp63β sites (Fig. 4C). Shared binding events are also significantly closer to either ΔNp63ɑ-specific (Fig. 4D) or ΔNp63β-specific (Fig. 4E) gene targets than isoform-specific binding events. ΔNp63ɑ-specific binding events are significantly closer to unique ΔNp63ɑ gene targets than ΔNp63β-specific binding events (Fig. 4D). This statistical significance is not preserved between ΔNp63β-specific genes and binding events. Our data suggest that isoform-specific binding is likely only a minor contributor to differential gene expression driven by ΔNp63ɑ and ΔNp63β, and that binding sites that are shared between ΔNp63ɑ and ΔNp63β are associated with both shared and isoform-specific gene regulation.

### ΔNp63β binding correlates with increased H3K27ac relative to ΔNp63ɑ

ΔNp63ɑ and ΔNp63β bind to highly similar, although not identical, genomic locations. Differential binding is associated with minor variations in the enrichment of transcription factor motifs. ΔNp63β has a stronger preference for canonical p53 family motifs and ΔNp63ɑ binding sites are more enriched with canonical AP-1 family motifs common in regulatory elements (Fig. 3D-E). ΔNp63ɑ activity is a pioneer factor and is involved in establishment and maintenance of epithelial-specific enhancers (Fessing et al., 2011; Bao et al., 2015; Karsli Uzunbas et al., 2019; Li et al., 2019). Isoform-specific binding is only weakly correlated with differential gene expression, as the majority of specific transcriptional differences are associated with common binding events (Fig. 4C-E). We therefore examined whether regulatory element activity at unique and shared p63 binding sites might correlate with the observed differences in gene expression between ΔNp63ɑ and ΔNp63β. We performed biological replicate ChIP-seq experiments for the histone modification H3K27ac, an established proxy for regulatory element activity, in HCT116 *TP53^-/-^* cell lines expressing either a negative control, ΔNp63ɑ, or ΔNp63β. These cells do not endogenously express p53, a strong, constitutive activator, which can influence H3K27ac and transcriptional dynamics at p63 binding sites. Thus, any changes in local chromatin should reflect local p63 isoform activity.

We observe high concordance in H3K27ac enriched regions (peaks) across control, ΔNp63ɑ and ΔNp63β-induced conditions (Fig. 5A). Less than 15% of either ΔNp63ɑ or ΔNp63β binding events overlap an H3K27ac peak in the negative control cell line (Fig. 5B-C). This is substantially lower than the greater than 75% overlap between ΔNp63ɑ binding sites and H3K27ac observed in the basal epithelial cell line MCF10A (Karsli Uzunbas et al., 2019). The percentage of p63 sites intersecting H3K27ac increases slightly in ΔNp63ɑ and ΔNp63β-induced conditions (Fig. 5B-C), suggesting that binding of these isoforms might be related to changes in H3K27ac enrichment. We next examined the intersection of p63 isoforms with H3K27ac peaks shared or uniquely enriched in isoform-specific cell lines. Approximately 12% of H3K27ac peaks found across control, ΔNp63ɑ, and ΔNp63β-induced conditions are bound by either ΔNp63ɑ (Fig. 5D) or ΔNp63β (Fig. 5E), and this co-occupancy drops dramatically at H3K27ac peaks found in common across control and either ΔNp63ɑ or ΔNp63β conditions. ΔNp63ɑ-specific H3K27ac peaks are more likely to be occupied by ΔNp63ɑ (12%) than those H3K27ac peaks found in p63-deficient conditions (4.3%)(Fig. 5D). Strikingly, we observe a near 10-fold increase (33% vs. 3.5%) in H3K27ac peaks found uniquely after ΔNp63β induction that are occupied by ΔNp63β relative to control H3K27ac (Fig. 5E). This increase in co-occupancy is similar at H3K27ac enriched regions found in ΔNp63ɑ and ΔNp63β conditions, but not in control (Fig. 5E). We then examined H3K27ac dynamics at either ΔNp63ɑ or ΔNp63β binding sites by comparing H3K27ac enrichment in isoform-specific conditions relative to negative controls. H3K27ac enrichment increases at least 2-fold at 556 ΔNp63ɑ binding sites (Fig. 5F) and at 1,851 ΔNp63β sites (Fig. 5G), whereas loss of H3K27ac after p63 binding is virtually non-existent. Although ΔNp63ɑ binding sites see an increase in H3K27ac enrichment, the gain in H3K27ac is more pronounced at ΔNp63β binding sites, consistent with our peak-based analysis (Fig. 5D-E).

**Figure 5:**
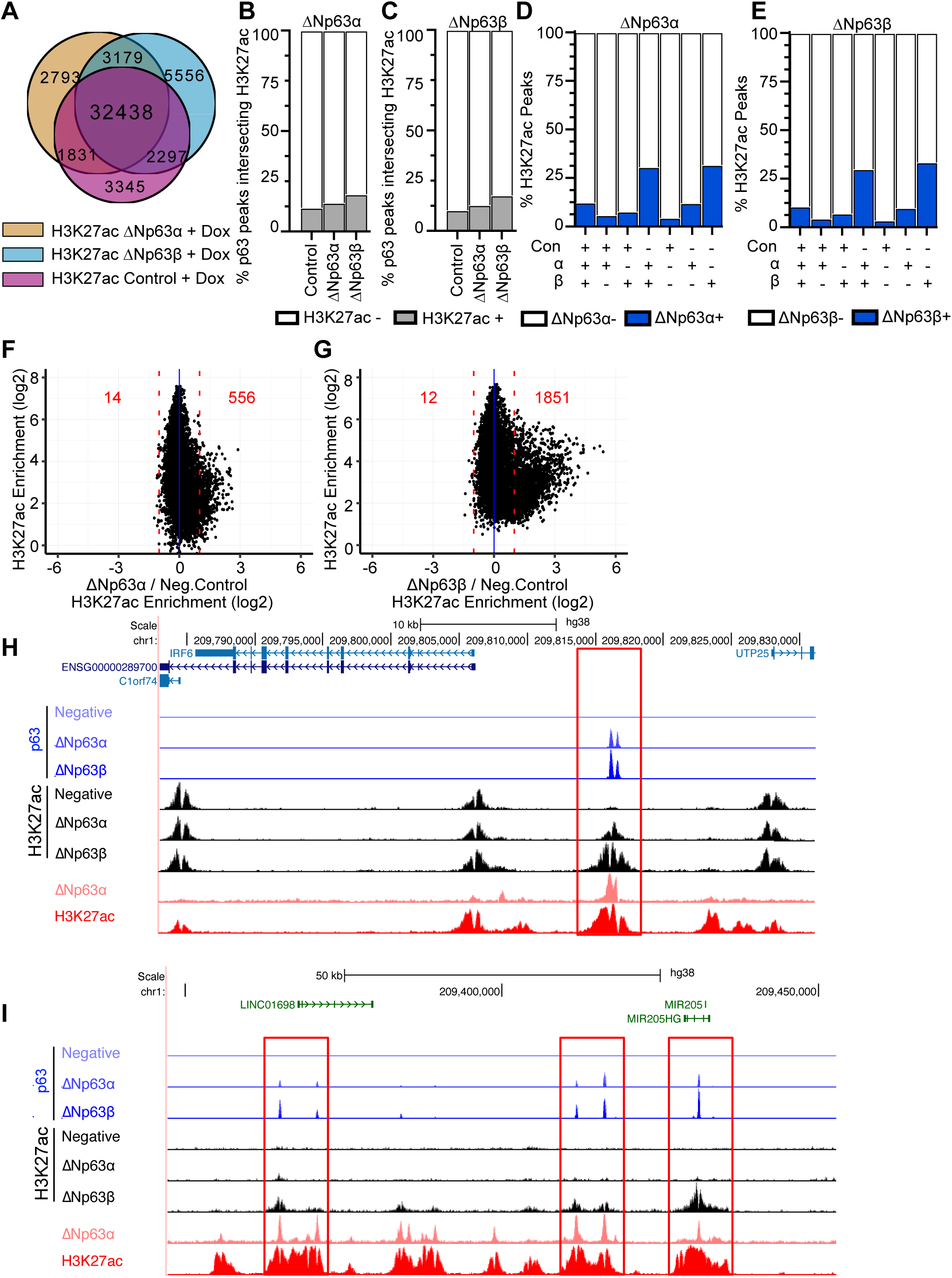
Regulatory element-associated H3K27ac dynamics at ΔNp63ɑ and ΔNp63β binding sites. **A)** The percentage of either ΔNp63ɑ or **B)** ΔNp63β ChIP-seq peaks intersecting MACS2-derived H3K27ac peaks from control, ΔNp63ɑ, or ΔNp63β induced conditions. **C)** Overlap of H3K27ac peaks found in control, ΔNp63ɑ, or ΔNp63β induction conditions. Biological replicates for each condition were first merged and then intersected using bedTools. **D)** The percent of H3K27ac peaks from each category shown in (C) with either D) ΔNp63ɑ binding sites or **E)** ΔNp63β binding sites. **F)** H3K27ac enrichment (log2 normalized) dynamics between ΔNp63ɑ or **G)** ΔNp63β conditions and negative control conditions. **H)** Genome browser view of *IRF6* locus and the **I)** MIR205 locus displaying p63 and H3K27ac ChIP-seq enrichment data for control, ΔNp63ɑ, or ΔNp63β-induced cell lines. The bottom two tracks represent p63 or H3K27ac ChIP-seq data from MCF10A mammary epithelial cell lines.

We then investigated H3K27ac and p63 binding dynamics by examining specific genomic loci near known target genes. The *IRF6* gene is activated by both ΔNp63ɑ or ΔNp63β. *IRF6 is* regulated by an upstream enhancer bound by p63, and loss of p63-dependent enhancer activity is associated with epithelial dysfunction and cleft palate in humans and mice (Rahimov et al., 2008; Thomason et al., 2010). Both ΔNp63ɑ and ΔNp63β bind to this upstream enhancer element and we observe a strong, binding site-specific gain in H3K27ac relative to negative control conditions (red box, Fig. 5H). This gain in H3K27ac only after p63 binding suggests a p63-dependent increase in *IRF6* enhancer activity that is associated with *IRF6* expression, and that this ability is shared by both isoforms. ΔNp63β, and not ΔNp63ɑ, uniquely induces expression of the epithelial-specific microRNA *MIR205* in HCT116 *TP53^-/-^* even though both p63 isoforms are capable of binding to nearby regulatory elements (Fig. 5I). Interestingly, only ΔNp63β binding is associated with increased H3K27ac enrichment at epithelial-specific regulatory regions (MCF10A H3K27ac, bottom, Fig. 5I). Occupancy of ΔNp63ɑ and ΔNp63β regulatory elements linked to other known p63-regulated genes *S100A2* (Fig. S2A), *ZNF750* (Fig. S2B), and *SFN* (Fig. S2C) are not associated with dynamic H3K27ac enrichment, suggesting that gains in H3K27ac at p63-bound regulatory elements are not strictly required for p63-dependent gene regulation. Taken together, our analysis of H3K27ac dynamics at p63 binding sites suggests a shared ability of ΔNp63ɑ and ΔNp63β to regulate local H3K27ac dynamics at gene regulatory elements, but that ΔNp63β unique relationship with H3K27ac enrichment may relate to its increased number of gene regulation targets.

### The TAD2 and Δ5 domains are critical for high transcriptional activity and target gene expression of ΔNp63β

ΔNp63ɑ and ΔNp63β share a set of gene targets, have highly similar genomic occupancy, and can both increase regulatory element activity, but the mechanisms that confer differential gene regulation are not clear. ΔNp63β has higher transcriptional activity in reporter assays, regulates a larger number of gene targets, and its genomic binding is associated with novel gains in H3K27ac at gene regulatory elements. Because their genomic binding profiles were highly similar (Fig. 3A) and were not strongly associated with differences in gene expression (Fig. 4C-E) , we reasoned that differences between ΔNp63ɑ and ΔNp63β likely lie in unique C-terminal domains. ΔNp63ɑ and ΔNp63β both share a second transactivation domain, or “TAD2”, located from AA 356-456, directly after the oligomerization domain. ΔNp63β also has five unique, C-terminal amino acids “Δ5” (AA 457-461) and lacks the SAM and ID domain found in ΔNp63ɑ. To determine the extent to which unique and shared domains contribute to p63β function, we created a series of C-terminal mutants in both ΔNp63β and TAp63β and tested their ability to activate transcription.

We cloned ΔTAD2, which removed the entire C-terminal region after the oligomerization domain (Fig. 6A) and Δ5, which removes the p63β-specific 5 amino acids at the C-terminus (Fig. 6A) and demonstrated expression in HCT116 *TP53^-/-^* cells (Fig. 6B). We then tested their ability to activate transcription of reporter (nanoLuciferase) downstream of a synthetic p63-response element derived from a regulatory element controlling the *SFN* gene (Hermeking et al., 1997). Deletion of either TAD2 or the Δ5 regions in TAp63β does not reduce transcriptional activity, and we observe a minor increase in the Δ5 mutant (Fig. 6C). We cannot rule out that this gain in transcriptional activity is due to increased expression of TAp63βΔ5 relative to wild-type TAp63β (Fig. 6B), but clearly, neither TAD2 or the β-specific 5AA C-terminus are required for transactivation by TAp63β (Fig. 6C). However, we observed clear requirements for these two domains for ΔNp63β activity. Deletion of the TAD2 region of ΔNp63β eliminates nearly all transcriptional activity (Fig. 6D), while removing β-specific 5AA domain significantly reduces the ability of ΔNp63β to activate this reporter (Fig. 6D). These data indicate the unique requirement of the C-terminal domains within ΔNp63β, but not TAp63β for transcriptional activation.

**Figure 6:**
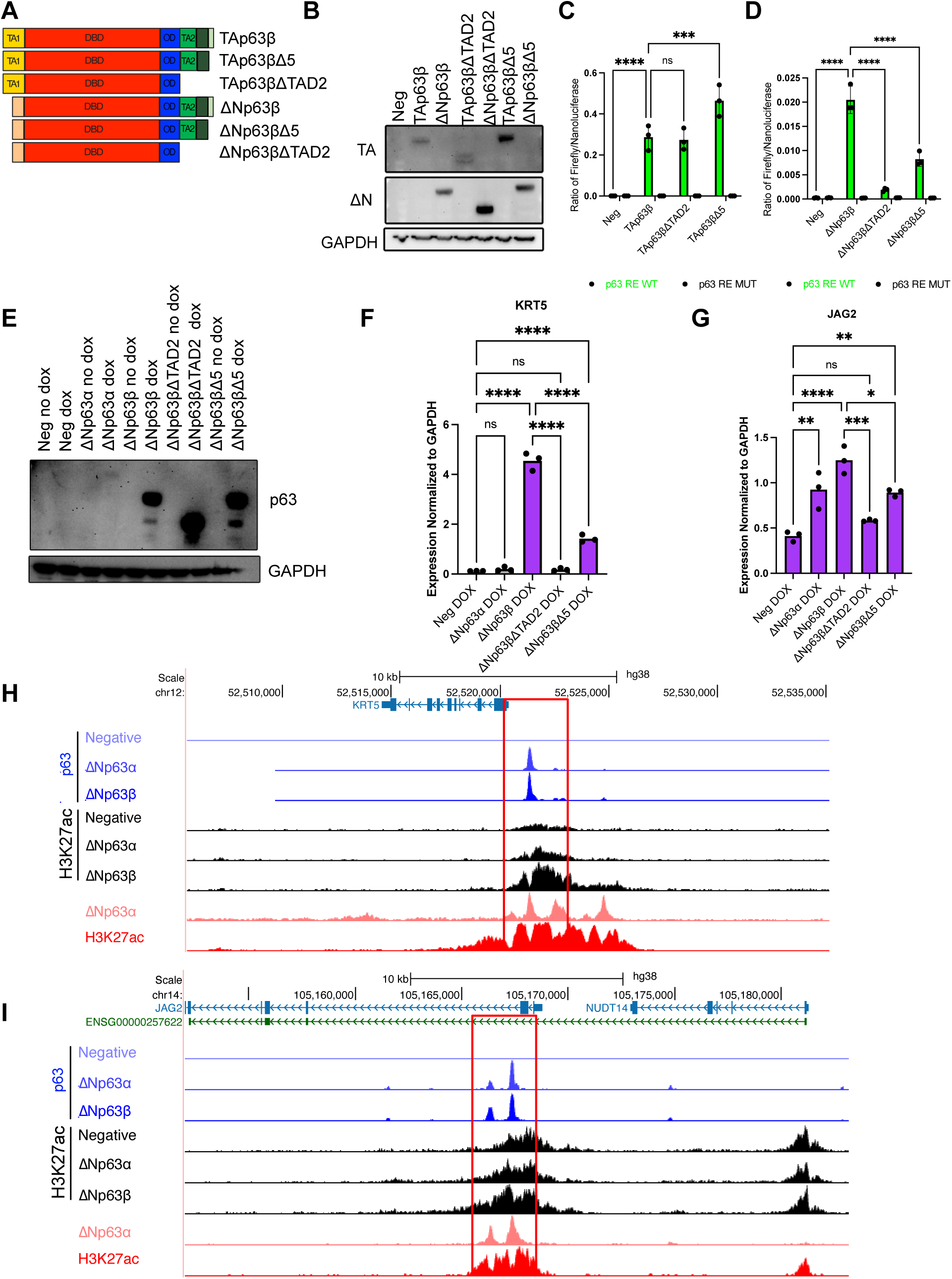
Analysis of C-terminal domain on ΔNp63β function. **A)** Schematic of TAp63 and ΔNp63 C-terminal TAD mutants. **B)** Protein expression of C-terminal TAD mutants in pcDNA plasmid constructs transiently transfected in HCT116 *TP53^-/-^* cells. Negative control is an empty pcDNA backbone. **C)** Reporter assay of TAp63 and **D)** ΔNp63 C-terminal TAD mutants on a p63 RE (green) and a mutant (grey) p63 RE. **E)** Protein expression of ΔNp63 C-terminal TAD mutants in lentiviral vectors under 24hr doxycycline induction in HCT116 *TP53^-/-^* cells. Negative control is GUS expressed in pCW57.1 vector. **F)** QRT-PCR analysis of *KRT5* expression by ΔNp63 C-terminal TAD mutants **G)** QRT-PCR analysis of *JAG2* expression by ΔNp63 C-terminal TAD mutants. **H)** Genome browser view of *KRT5* locus displaying p63 and H3K27ac ChIP-seq binding data. **I)** Genome browser view of *JAG2* locus displaying p63 and H3K27ac ChIP-seq binding data. Statistical analysis for qRT-PCR data was done using a a One-way ANOVA test (*: *p*-value <.05, **: *p*-value <.01, ***: *p*-value <.001, ****: *p*-value < 0.0001, ns = not significant) and a Two-way ANOVA ( ***: *p*-value <.001, ****: *p*-value < 0.0001, ns = not significant ) for reporter assay data.

We then sought to determine if these C-terminal domains are required for ΔNp63β activity to regulate native target genes and not only an artificial reporter system. To this end, we created HCT116 *TP53^-/-^* cell lines expressing either WT ΔNp63β, ΔNp63βΔTAD2, or ΔNp63βΔ5 (Fig. 6E) and measured expression of either shared ΔNp63 genes or ΔNp63β-specific targets. ΔNp63β lacking TAD2 does not activate expression of ΔNp63β-specific target genes *KRT5* (Fig. 6F), *MDM2, MIR205HG, SNAI2* or *IL1A* (Fig. S3A-D). *KRT5* is specifically activated by ΔNp63β despite similar ΔNp63ɑ and H3K27ac enrichment (Fig. 6H). The ability of ΔNp63βΔ5 mutant to activate these target genes is reduced relative to wild-type (Fig. 6F, Fig. S3A-D), indicating the β-specific 5AA domain contributes to unique ΔNp63β activities. For target genes of both ΔNp63ɑ and ΔNp63β, we observe a similar trend where TAD2 and the β-specific 5AA are required for full transactivation by ΔNp63β (Fig. 6G). Full activation of *JAG2* (Fig. 6G) by ΔNp63β requires the β-specific 5AA, with ΔNp63βΔ5 displaying activity equivalent to ΔNp63ɑ which lacks this domain. TAD2, found in both ΔNp63ɑ and ΔNp63β, is required for ΔNp63β-dependent *JAG2* and *FAT2* expression (Fig. S4B). The ΔNp63βΔTAD2 mutant activates *FAT2* gene expression to about the same extent as ΔNp63ɑ, but substantially less than WT ΔNp63β, suggesting that TAD2 is not required for ΔNp63ɑ-dependent transactivation. Our results suggest that TAD2 and a β-specific 5AA C-terminal domain are critical for transcriptional activation by ΔNp63β and likely contribute to gene regulatory differences between ΔNp63 C-terminal isoforms.

### ΔNp63β contains a unique, β-specific TAD at its C-terminus

p63ɑ and p63β isoforms contain the TAD2 domain, located after the oligomerization domain from position 356-456 (relative to ΔN isoforms). We demonstrated this domain is critical for transcriptional activation of reporter genes and of native p63 targets by ΔNp63β. On the contrary, TAp63β activity is unaffected when the TAD2 domain is deleted (Fig. 6C). ΔNp63Δ contains a partial TAD2 domain (AA 356-408) and a unique C-terminal extension, but displays weak transactivation in reporter systems (Fig. 1F). We also noted the p63β-specific 5AA C-terminal domain is required for full transcriptional activation by ΔNp63β. To further explore biological activities conferred by the p63β-specific C-terminus, we expressed a series of C-terminal p63 variants (Fig. 7A-B) and tested their ability to activate transcription of a p63-dependent reporter in HCT116 *TP53^-/-^* cells.

**Figure 7:**
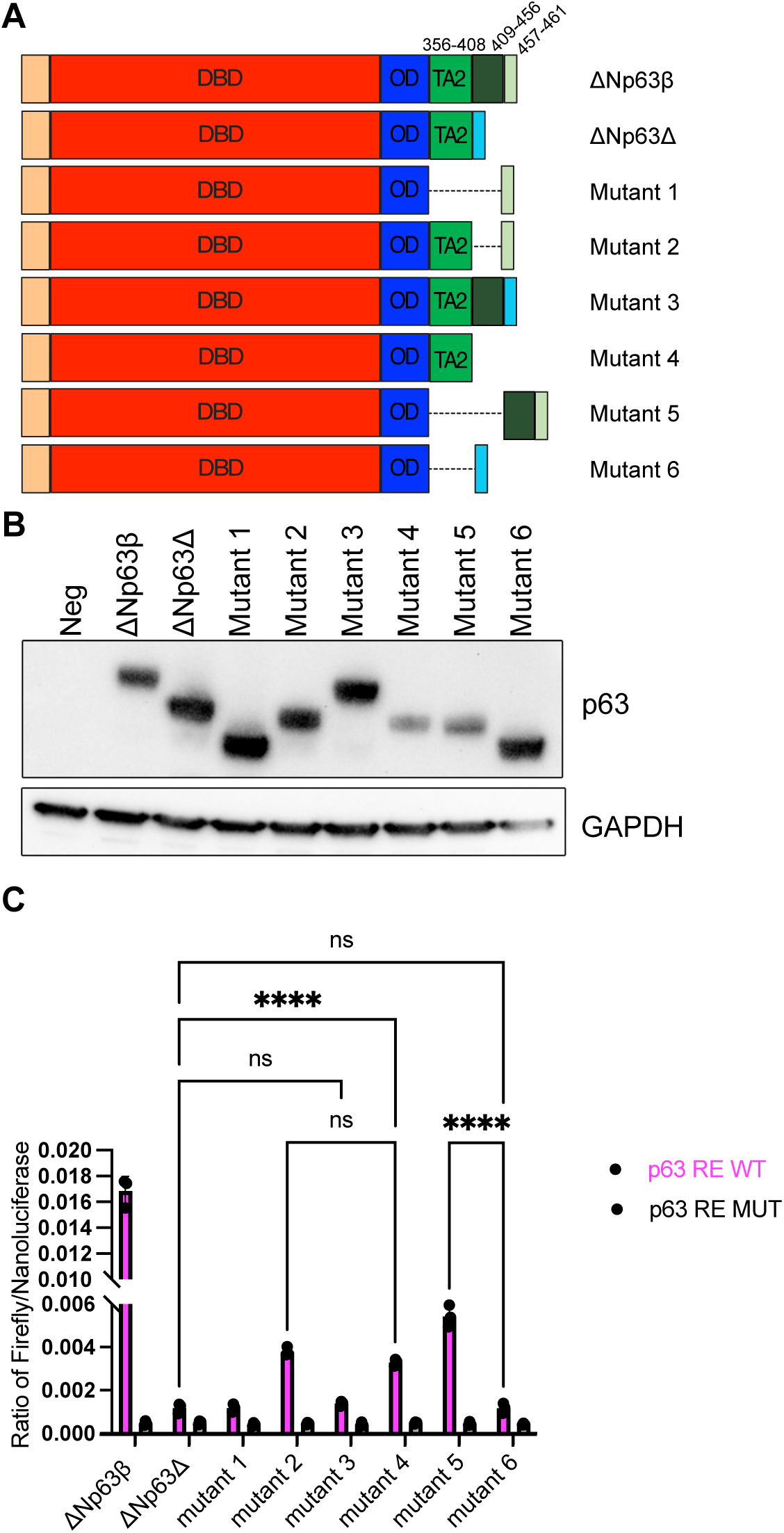
Characterization of function of ΔNp63 C-terminal isoform mutants. **A)** Schematic of ΔNp63 C-terminal mutants. **B)** Protein expression of C-terminal mutants transiently transfected in HCT116 *TP53^-/-^* cells expressed in a pcDNA backbone. Negative control is an empty pcDNA backbone. **C)** Reporter assay of p63 C-terminal mutants on using either a WT p63 RE (pink) or mutant p63 RE (grey). ΔNp63Δ and all C-terminal mutants have a statistically significant reduction in activity compared to ΔNp63β (****: *p*-value < 0.0001, ns = not significant, Two-way ANOVA)

We first asked whether the β-specific 5AA C-terminal domain might act as a third TAD, as it is required for full transactivation of ΔNp63β and is the only domain unique to ΔNp63β compared to ΔNp63ɑ. Mutant 1 removes TAD2 (AA356-456) from ΔNp63β, leaving the 5AA C-terminus directly next to the OD. Mutant 1 has weak activity when compared to WT ΔNp63β, and is comparable to ΔNp63Δ. These data suggest that the β-specific 5AA C-terminal domain is likely not an independent TAD. ΔNp63ɑ, ΔNp63β, and ΔNp63Δ share AAs 356-408 of TAD2, while AAs 409-456 are unique to ΔNp63ɑ and ΔNp63β (Fig. 7A). To determine the importance of these regions of TAD2 for ΔNp63β function, we created additional ΔNp63β variants which lack either AAs 409-456 (Mutant 2) or lack AAs 356-408 (Mutant 5), which is shared in ΔNp63ɑ, ΔNp63β, and ΔNp63Δ. Mutant 2 and Mutant 5 had comparable activity and displayed an approximately 3-fold decrease in transactivation compared to WT ΔNp63β (Fig. 7C). Importantly, both Mutant 2 and Mutant 5 are more active than either Mutant 1 or ΔNp63Δ, suggesting p63ɑ and p63β-specific AA 409-456 contributes to transcriptional activation.

ΔNp63Δ is less transactivating than Mutant 2, despite the only difference being the presence of unique C-terminal domains. ΔNp63Δ has eight unique amino acids on its C-terminus, compared to the 5AA specific to ΔNp63β. Removal of the 8 Δ-specific amino acids from ΔNp63Δ increases transactivation (Mutant 4) compared to WT ΔNp63Δ (Fig. 7C) suggesting these residues may repress transcription. This repressive effect of the ΔNp63Δ-specific 8AA C-terminus is supported by Mutant 3, where the Δ-specific domain is swapped for the 5AA β-specific domain. Mutant 3 activity is comparable to both ΔNp63Δ and Mutant 1, which lacks the entire TAD2 domain. Interestingly, removal of AA 356-408 from ΔNp63Δ (Mutant 6) is comparable to WT ΔNp63Δ further suggesting that the Δ-specific 8AA C-terminus likely represses the activity of TAD2 AA356-408. Our analysis of C-terminal variants of ΔNp63β suggests both AAs 356-408 and 409-456 are critical for full activity of ΔNp63β, but that Aas 409-456 require the presence of the 5AA β-specific C-terminus. Thus, while this β-specific 5AA C-terminus likely does not work independently in transcriptional activation, it appears to cooperate with 409-456 to form a unique, β-specific TAD.

## DISCUSSION

Genetic dissection of p63 activity has strongly implicated ΔNp63ɑ as essential for the establishment and maintenance of epithelial identity. However, the contribution of other p63 isoforms to these key biological activities remains largely unclear. Previous studies suggest that ΔNp63β can complement specific ΔNp63ɑ activities in vivo and possesses unique growth suppression abilities compared to ΔNp63ɑ. The specific mechanisms driving these behaviors, however, have not been fully explored. In this study, we analyze the genetic and molecular basis of differential gene expression networks driven by the C-terminal p63 isoforms ΔNp63ɑ and ΔNp63β. Our work confirms prior studies demonstrating that ΔNp63β has an increased ability to activate transcription driven by p63-responsive regulatory elements relative to ΔNp63ɑ and other C-terminal isoforms (Fig. 1F) (Helton et al., 2006). ΔNp63β contains a unique C-terminus relative to other isoforms that is required for transcriptional activation and control of a ΔNp63β-specific gene network (Fig. 6D,F. Fig. S3A-D). Although the functional impact of this increased transcriptional activation potential of ΔNp63β is not yet resolved, our data suggest key molecular and biochemical events that may provide clues into the observed differences between isoforms.

ΔNp63ɑ and ΔNp63β regulate a shared set of gene targets canonically associated with p63 activity, such as genes involved in epidermis development, tissue morphogenesis, and control of apoptosis (Fig. 2E). Despite higher activity in reporter assays (Fig. 2D), we did not observe universally higher RNA induction by ΔNp63β for these shared target genes. Both ΔNp63ɑ and ΔNp63β regulate specific gene networks (Fig. 2D,G). They bind to numerous shared genomic loci, and these shared sites are more closely associated with p63-induced gene expression than sites bound preferentially by a single isoform (Fig. 4C-E). This is true even for genes uniquely controlled by either isoform (Fig. 4D-E). The genomic location and occupancy of isoform-specific binding events suggest that isoform-specific binding plays only a modest role in differential transcriptional activity (Fig. 4D-E) relative to shared binding events. Thus, differential genomic occupancy of ΔNp63ɑ and ΔNp63β is unlikely to explain most isoform-specific gene regulatory events. This suggests that context-dependent, isoform-specific activity at shared gene regulatory elements may control differential gene expression potential.

Binding of ΔNp63ɑ or ΔNp63β to the genome correlates with increased enrichment of H3K27ac, a hallmark of regulatory element activity. We assayed p63 binding and H3K27ac enrichment in HCT116 *TP53^-/-^* cell lines lacking endogenous expression of both p53 and p63, allowing the analysis of p63 isoform-specific gene regulation, genomic binding, and chromatin dynamics. Increased enrichment of H3K27ac is more pronounced and widespread for ΔNp63β, with numerous ΔNp63ɑ and ΔNp63β binding sites gaining H3K27ac enrichment only after binding by ΔNp63β. Therefore, one mechanism underlying differential gene expression by ΔNp63ɑ and ΔNp63β may be an increased in the ability of ΔNp63β to effect changes in local chromatin structure at gene regulatory elements. The specific molecular mechanisms underlying p63-dependent regulation of local and long-distance chromatin structure, including H3K27ac deposition, are not yet fully known (Kouwenhoven et al., 2015; Li et al., 2019; Lin-Shiao et al., 2019). ΔNp63ɑ directly interacts with HDAC1 and HDAC2 via the ɑ-specific TID which may contribute to lower local H3K27ac (LeBoeuf et al., 2010; Ramsey et al., 2011). Both ΔNp63ɑ and ΔNp63β interact with the acetyl-binding and transcriptional co-activator protein BRD4 to regulate keratinocyte-specific gene expression (Foffi et al., 2024). Ultimately, a deeper investigation into the shared and isoform-specific molecular mechanisms of gene regulation is required better understand the context-dependent differences and biological activities of ΔNp63ɑ and ΔNp63β.

Our data suggest that the increased transcriptional activity of ΔNp63β relative to ΔN isoforms is likely due to the second transactivation domain (TAD2) and the β-specific inclusion of a uniquely activating C-terminal domain. The five amino acid, β-specific C-terminus is necessary for full transcriptional activation by ΔNp63β (Fig. 6D). Recent work suggests a short, β isoform-specific domain is crucial for activity of the p63β paralog p73β (Li et al., 2024). This five amino acid, C-terminal domain in p73β was necessary for both TAp73β and ΔNp73β, whereas our results suggest this domain may be dispensable for transcriptional activation of TAp63β. Both studies suggest the short β-specific domain of p73 and p63 works in conjunction with amino acids in a domain directly upstream (Fig. 7B-C). Although loss of the β-specific domain reduces transcriptional activity, replacement of this domain with the short, p63Δ-specific C-terminus completely ablates transcriptional activity, and removal of this Δ-specific C-terminus from ΔNp63Δ significantly increases transactivation ability. Thus, it appears the Δ-specific C-terminal domain may confer unique, transcriptional repression properties on ΔNp63Δ. TAp63β,γ, and Δ are all strongly transactivating compared to TAp63ɑ, whereas only ΔNp63β displays high transactivation potential across the ΔN isoforms (Fig. 1E-F). These observations suggest p63 C-terminal splice variants may have differential effects on TA and ΔN isoforms, like for p63ɑ. The ɑ-specific SAM and TI domains inhibit TAp63ɑ by adopting a unique inhibitory conformation, but independently repress ΔNp63ɑ through interactions with co-repressor proteins or via ubiquitin-mediated degradation (Ghioni et al., 2002; Serber et al., 2002; Li et al., 2008; Pecorari et al., 2022). Our results provide evidence that the broadly expressed p63 C-terminal variant ΔNp63β controls a unique gene regulatory network compared to ΔNp63ɑ through a β-specific C-terminus. How different C-terminal splice variants, and their included or excluded protein domains, elicit unique biological activities across p63 isoforms remains an open question. These and other recent data provide further evidence for p63 isoform-specific biological function, and future work should focus on resolving the spatial and temporal context for these differential activities.

## MATERIALS AND METHODS

### Cell Culture

HCT116 *TP53^-/-^* cells were cultured in McCoys media (Gibco, #16-600-082) supplemented with 10% FBS (Corning, #35-016-CV) and 1% penicillin-streptomycin (Gibco, #15240-062). Human embryonic kidney cell line HEK293FT were cultured in Dulbecco’s Modified Eagle’s Medium 1X (Corning 10-013-CV) and supplemented with 10% FBS and 1% penicillin-streptomycin. For doxycycline inducible cell lines, doxycycline was added at 500ng/ml 24 hours before collection. All cell lines were cultured at 37°C and 5% CO_2_.

### Plasmids and Cloning

p63 isoform plasmids were originally obtained from Twist Biosciences, whereby they were either cloned into pcDNA3.1 mammalian expression vector for transient expression or pCW57.1 lentiviral vector for integrated, doxycycline inducible expression. GUS control plasmid was provided as part of the LR Clonase II enzyme kit (Invitrogen 11791020). Due to the design of the Twist plasmids, AgeI sites were cloned into the pcDNA3.1 MCS via site-directed mutagenesis, and p63 isoforms were cloned by restriction digest of AgeI and BglII (BamHI) sites and ligation into pcDNA3.1. For pCW57.1, p63 isoforms in pENTR Twist backbone (Twist Biosciences) were cloned via Gateway cloning using LR Clonase enzyme. All mutants were cloned via site directed-mutagenesis or HiFi assembly and full plasmid sequencing was performed using Plasmidsaurus. All primers and plasmid information are listed in Table S1.

### Lentiviral Production

HEK293FT cells were seeded at a density of 600,000 cells in a 6-well plate. One microgram of pCW57.1 lentiviral plasmid was transfected along with 600ng psPAX2 and 400ng pMD2.G (pCW57.1 was a gift from David Root, Addgene plasmid # 41393 ; http://n2t.net/addgene:41393; RRID:Addgene_41393), psPAX2, and pMD2.G (psPAX2 and pMD2.G were a gift from Didier Trono, Addgene plasmid # 12260 ; http://n2t.net/addgene:12260; RRID:Addgene_12260) were obtained from Addgene). Lentiviral supernatant was collected at 24 and 48 hours. Cell lines to be infected were seeded at a density of 400,000 and infected with viral supernatant that was concentrated using spin dialysis, along with 8ug/ml polybrene. Viral supernatant was removed from cells after 24 hours and replaced with fresh media. Forty-eight hours after infection, cell lines infected with pCW57.1 vectors were selected with 2ug/ml puromycin for 72 hours.

### Western Blotting

Protein was isolated using custom made RIPA buffer (50 mM Tris-HCl pH 7.4, 150 mM NaCl, 1% NP-40, 0.5% sodium deoxycholate, 0.1% SDS, 1mM EDTA, 1% Triton x-100) supplemented with protease inhibitor (Pierce, 78442). Concentration of isolated protein was measured using a microBCA kit (Pierce, 23227) and 25µg was loaded on a 4-12% Bis-Tris protein gel (Invitrogen, NP0321BOX). Protein size was analyzed using PageRuler™ Prestained Protein Ladder (Thermo 26616). Membranes were blocked in 5% non-fat milk in TBS-T. Antibodies used included rabbit anti-ΔNp63 antibody (Cell Signaling E6Q3O), mouse anti-TAp63 (BioLegend 938102), rabbit anti-p63 DBD (abcam97865), and rabbit anti-GAPDH antibody (Cell Signaling D16H11).

### Reporter assays

The BDS-2,3 p63 responsive element from the *SFN* gene was cloned into the pGL4.24 vector (Hermeking et al., 1997). Luciferase assays were carried out using Nano-Glo® Dual-Luciferase® Reporter Assay System (Promega #1620). HCT116 *TP53^-/-^* cells were seeded at a density of 50,000 cells in a 96-well plate and transfected via reverse transfection. PGL4.24 firefly vector (GenBank® Accession Number: DQ904456) was used as reporter backbone and pNL1.1 nanoluciferase, with constitutive PGK promoter, (Promega #N1441) was used as a normalizing control vector. p63 isoforms and isoform mutants cloned into the pcDNA3.1 vector were transiently reverse transfected alongside reporter gene constructs (Polyplus #101000046) at a concentration of 200ng for isoform constructs and 180ng for luciferase constructs.

### RNA isolation and RT-qPCR

RNA isolation was carried out using Quick RNA Miniprep kit (Zymo, #R1055) and cDNA was generated (Thermo 4368813). qPCR was performed using iTaq Universal SYBR Green Supermix (Bio-Rad #1725121) and utilizing the relative standard curve method. qPCR primers are listed in Table S1.

### Gene Expression Analysis using RNA-seq

Doxycycline-inducible ΔNp63ɑ, ΔNp63β, or a negative control (Gus) HCT116 *TP53^-/-^* cells were generated using lentiviral transduction as described above. For each cell type, three biological replicates for each isoform or control cell line were seeded at a density of 400,000 cells. The day after seeding, doxycycline (500ng/mL) was added to induce protein expression. Twenty-four hours after induction cell pellets were collected and RNA isolation was carried out as described above. RNA-seq compatible libraries were constructed after polyA-selection and sequenced on an Illumina HiSeq 2000 by Azenta. Reads were quantified using *kallisto* in bootstrap mode (n=100) against the Ensembl transcriptome (v. 104) and differentially-expressed genes were called using *DeSeq2* (Love et al., 2014; Bray et al., 2016).

### ChIP-seq of p63 and H3K27ac

ΔNp63ɑ, ΔNp63β and GUS negative control cell line were seeded and treated with 500ng/ml doxycycline for 24 hours. Twenty-five million HCT116 *TP53^-/-^*cells per replicate and two biological replicates were prepared using Diagenode iDeal ChIP-seq kit for Transcription Factors (Diagenode #C01010170). Samples were fixed with 1% formaldehyde for 10 minutes followed by quenching with 250mM glycine. Chromatin was sheared using the Diagenode Bioruptor Plus for 50 cycles (30s on/off). Antibodies used include anti-ΔNp63 antibody (Cell Signaling E6Q3O), and anti-H3K27ac antibody (Diagenode C15410196). DNA sequencing libraries were prepared using the NEBNext Ultra II DNA Library prep kit using standard protocols. Samples were sequenced using a NextSeq 2000 (2x50bp) at the University at Albany Center for Functional Genomics. Paired-end sequencing reads were aligned to the human hg38 reference genome using HiSat2 (Kim et al., 2019) with unaligned reads omitted from the resulting output (--no-unal). Aligned reads were sorted by position and converted to *bam* format using *samtools* (Li et al., 2009). Regions of enrichment (peaks, q-value <= 0.01) for p63 were called with p63 ChIP-seq from empty vector-expressing HCT116 *TP53^-/-^*cells as a background control using *macs2* (Zhang et al., 2008). H3K27ac peaks were called without background controls. Peaks within problematic genomic regions were removed based on the ENCODE blacklist using *bedtools* (Amemiya et al., 2019). Peak intersection analysis was performed using *bedtools* with any overlap considered a positive intersection, and peaks from biological replicates were merged to create a high-confidence peak set before downstream analysis and comparison with additional datasets (Quinlan and Hall, 2010). Venn diagrams for peak intersection analysis were generated using *intervene* (Khan and Mathelier, 2017). Heatmaps, bigwig files, and quantification of read enrichment within regions of interest were generated using *deeptools2* (Ramírez et al., 2016). Gene ontology analysis was performed using *metascape* on a local Docker installation (Zhou et al., 2019). Complete Gene Ontology (GO) analysis can be found in Table S2.

### ChIP-seq, motif enrichment, and nearest gene analysis

Motif enrichment within p63 peak regions was performed using a size and GC-matched genomic background using findMotifsGenome script from *HOMER* (Heinz et al., 2010). p53 family motifs in the hg38 reference genome were identified using JASPAR motif models (p53: MA0106.3, p63: MA0525.2, p73: MA0861.1) identified using scanMotifGenomeWide package in *HOMER* and then merged with p63 peak locations (Castro-Mondragon et al., 2022). Genes or transcriptional start sites nearest to p63 binding sites were identified using closestBed from the *R-*implementation of *bedtools* (*bedtoolsr, v.2.30.0-5*) and statistics were calculated using the *rstatix* package (0.7.2). The *polyEnrich* module of the *R* implementation of *chipEnrich* (v.2.26.0) was used to examine Gene Ontology of TSS nearest p63 binding sites (Welch et al., 2014).

## DATA AVAILABILITY

All sequencing datasets are available through Gene Expression Omnibus (GSE283357 for RNA-seq and GSE283359 for ChIP-seq).

## ACKNOWLEDGMENTS

This work was supported by a grant from the National Institutes of Health (NIGMS R35138120) to MAS. The authors would like to acknowledge support from the University at Albany Center for Functional Genomics.

---

**Supplementary Figure 1:**
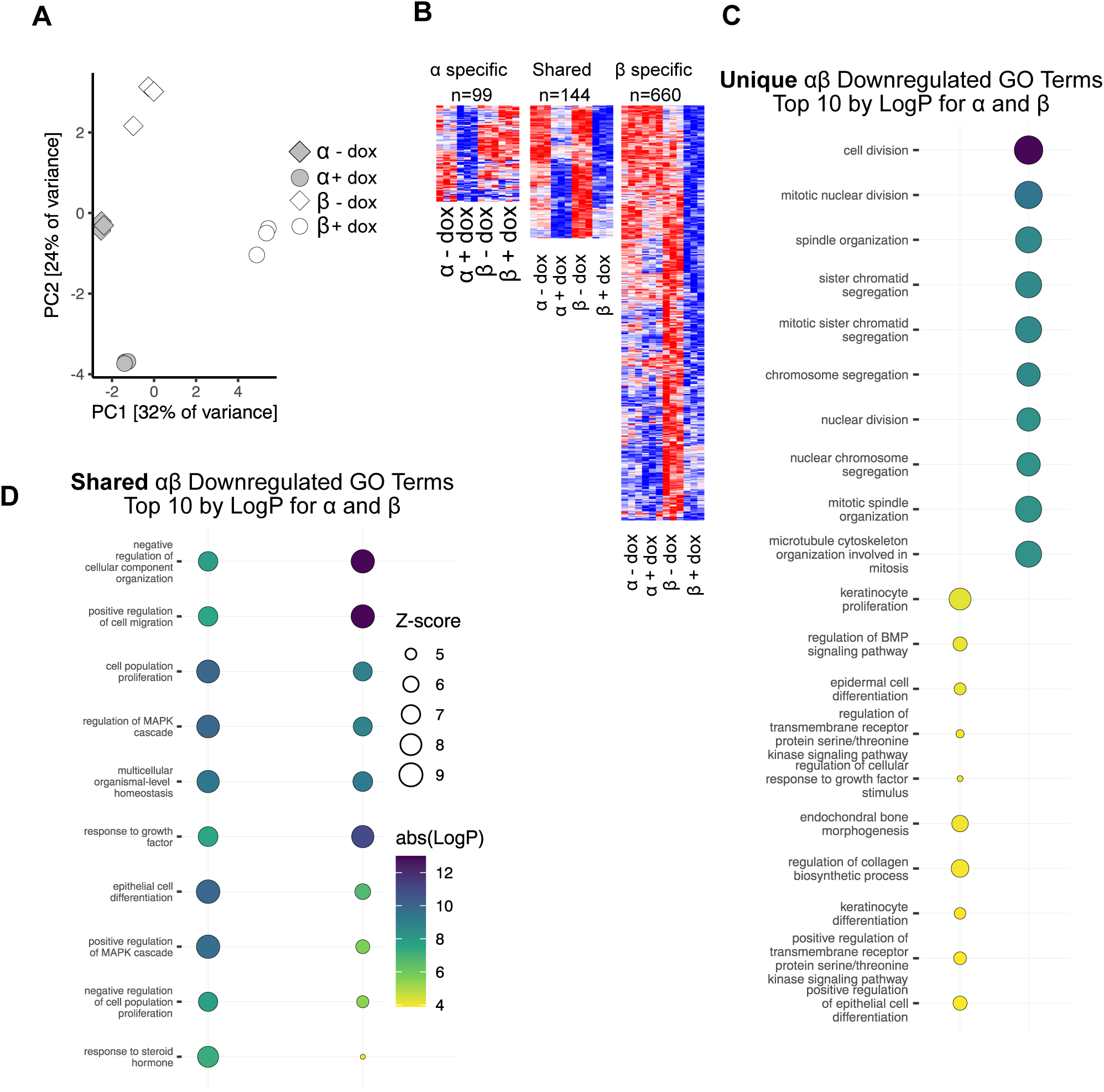
RNA-seq analysis of shared and unique downregulated genes. Principal component analysis (PCA) plot for the 4,000 most highly-expressed genes from RNA-seq analysis of either ΔNp63ɑ or ΔNp63β induction (-/+ 500ng/mL doxycycline for 24hrs) in HCT116 *TP53^-/-^* cells. **B)** Heatmaps of differentially downregulated genes (q-value <= 0.05) that are shared, or specific to either ΔNp63ɑ or ΔNp63β-induced conditions. **C)** Gene ontology enrichment of downregulated genes after induction of either ΔNp63ɑ or ΔNp63β, showing the Top 10 Gene Ontology terms uniquely identified in either ΔNp63ɑ or ΔNp63β differentially downregulated genes. **D)** Top 10 Gene Ontology terms for shared downregulated genes after induction of either ΔNp63ɑ or ΔNp63β.

**Supplementary Figure 2:**
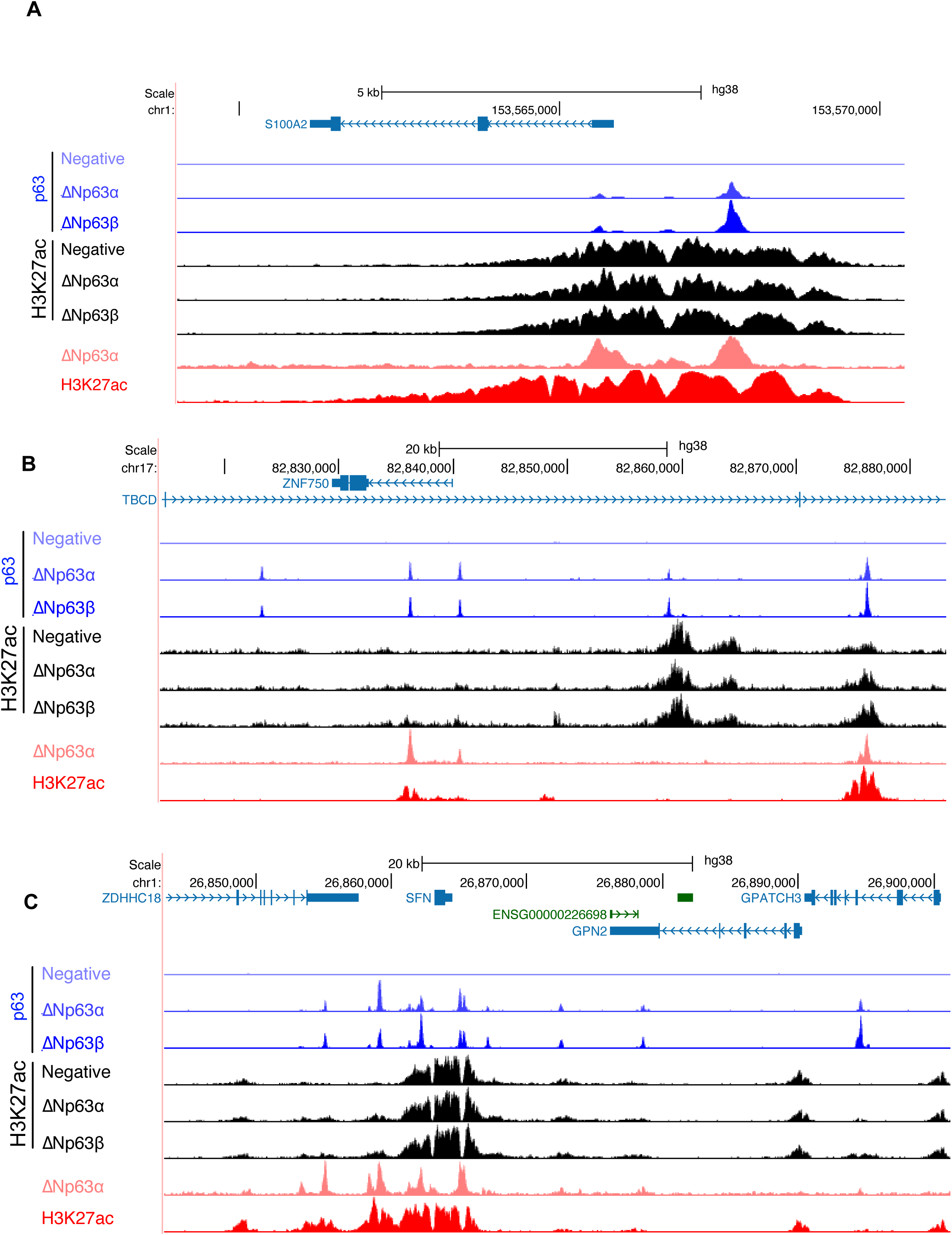
ChIP-seq analysis of p63 and H3K27ac at genomic loci. UCSC Genome browser view of ChIP-seq of control, ΔNp63ɑ, or ΔNp63β and H3K27ac at **A)** *S100A2* **B**) *ZNF750* and **C)** *SFN* loci.

**Supplementary Figure 3:**
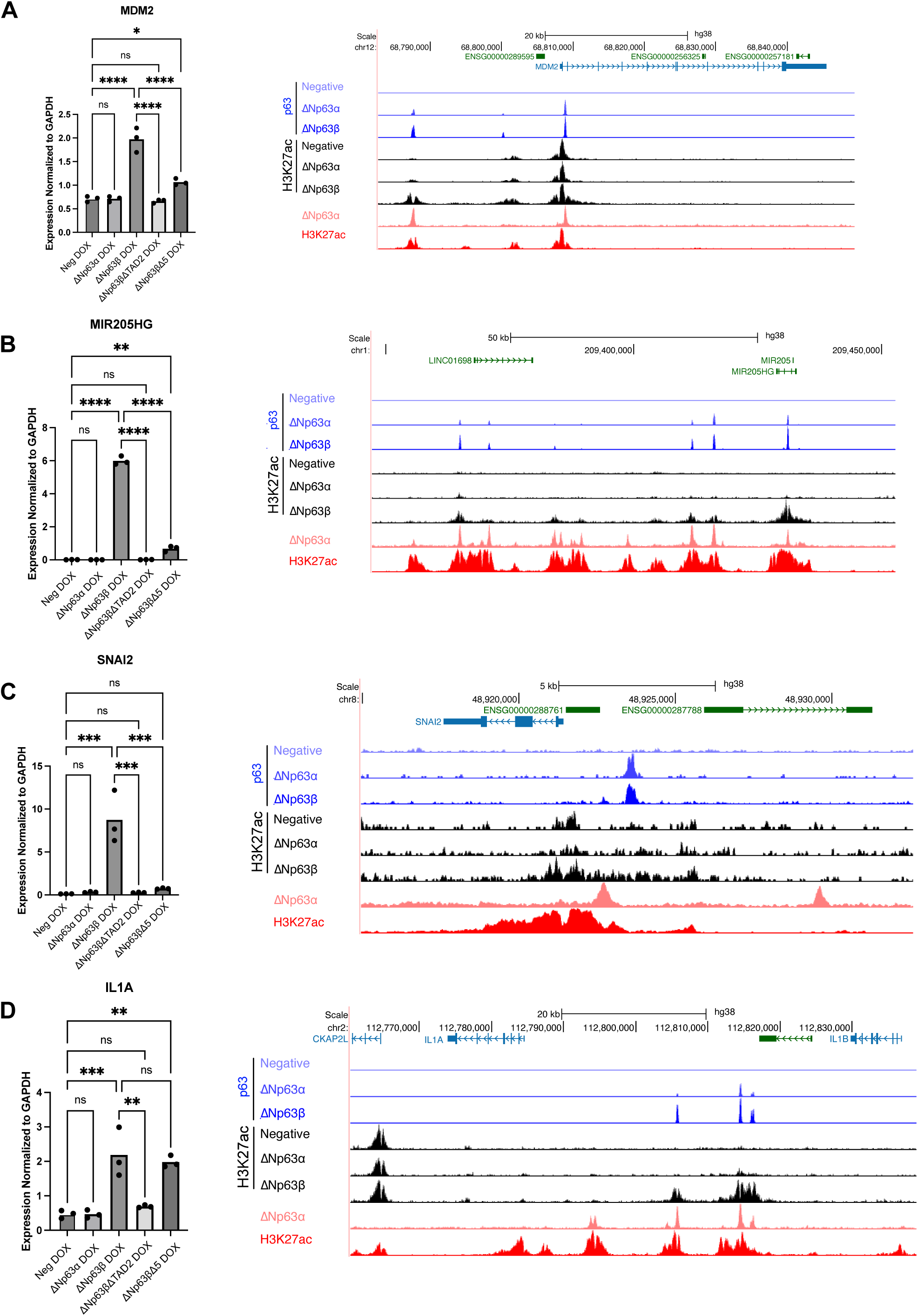
Analysis of C-terminal domain on ΔNp63β function at ΔNp63β-only upregulated genes. qRT-PCR analysis of **A)** *MDM2* **B)** *MIR205HG* **C)** *SNAI2* and **D)** *IL1A* expression by ΔNp63 C-terminal TAD mutants and UCSC genome browser views of their respective genomic loci, showing p63 and H3K27ac enrichment. (*: *p*-value <.05, **: *p*-value <.01, ***: *p*-value <.001, ****: *p*-value < 0.0001, ns = not significant, One-way ANOVA)

**Supplementary Figure 4:**
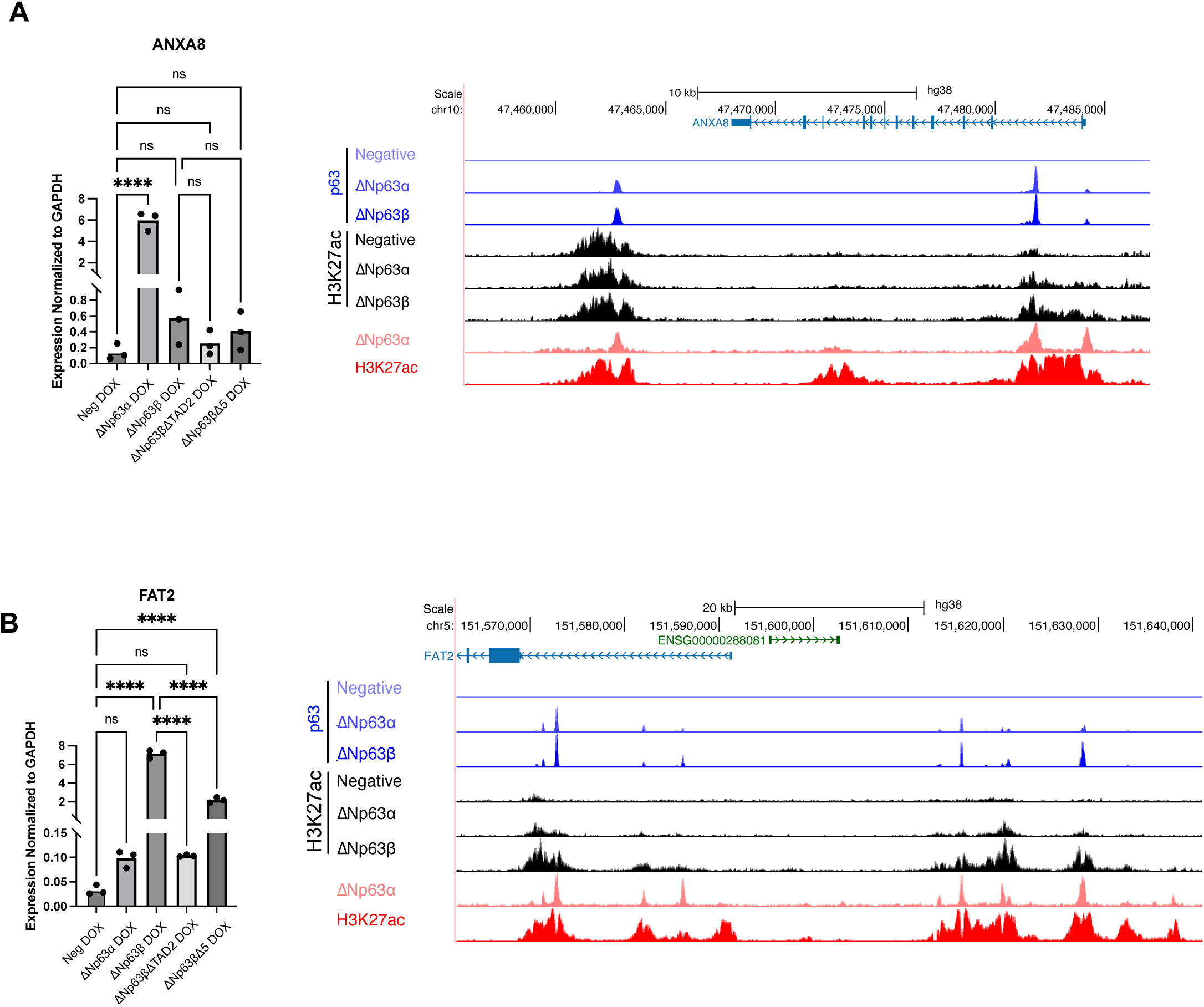
Analysis of C-terminal domain on ΔNp63β function at ΔNp63β and ΔNp63ɑ upregulated genes. qRT-PCR analysis of **A)** *ANXA8* and **B)** *FAT2* expression by ΔNp63 C-terminal TAD mutants and genome browser views of their respective genomic loci, showing p63 and H3K27ac enrichment. (****: *p*-value < 0.0001, ns = not significant, One-way ANOVA)

## WORKS CITED

1. Amemiya, H. M., Kundaje, A., and Boyle, A. P. (2019). The ENCODE Blacklist: Identification of Problematic Regions of the Genome. Sci Rep 9, 9354. doi: 10.1038/s41598-019-45839-z

2. Bao, X., Rubin, A. J., Qu, K., Zhang, J., Giresi, P. G., Chang, H. Y., et al. (2015). A novel ATAC-seq approach reveals lineage-specific reinforcement of the open chromatin landscape via cooperation between BAF and p63. Genome Biology 16, 284. doi: 10.1186/s13059-015-0840-9

3. Bray, N. L., Pimentel, H., Melsted, P., and Pachter, L. (2016). Near-optimal probabilistic RNA-seq quantification. Nat Biotechnol 34, 525–527. doi: 10.1038/nbt.3519

4. Castro-Mondragon, J. A., Riudavets-Puig, R., Rauluseviciute, I., Berhanu Lemma, R., Turchi, L., Blanc-Mathieu, R., et al. (2022). JASPAR 2022: the 9th release of the open-access database of transcription factor binding profiles. Nucleic Acids Research 50, D165– D173. doi: 10.1093/nar/gkab1113

5. Celli, J., Duijf, P., Hamel, B. C. J., Bamshad, M., Kramer, B., Smits, A. P. T., et al. (1999). Heterozygous Germline Mutations in the p53 Homolog p63 Are the Cause of EEC Syndrome. Cell 99, 143–153. doi: 10.1016/S0092-8674(00)81646-3

6. Coutandin, D., Osterburg, C., Srivastav, R. K., Sumyk, M., Kehrloesser, S., Gebel, J., et al. (2016). Quality control in oocytes by p63 is based on a spring-loaded activation mechanism on the molecular and cellular level. eLife 5, e13909. doi: 10.7554/eLife.13909

7. Di Girolamo, D., Di Iorio, E., and Missero, C. (2024). Molecular and Cellular Function of p63 in Skin Development and Genetic Diseases. Journal of Investigative Dermatology. doi: 10.1016/j.jid.2024.08.011

8. Dohn, M., Zhang, S., and Chen, X. (2001). p63α and ΔNp63α can induce cell cycle arrest and apoptosis and differentially regulate p53 target genes. Oncogene 20, 3193–3205. doi: 10.1038/sj.onc.1204427

9. Fessing, M. Y., Mardaryev, A. N., Gdula, M. R., Sharov, A. A., Sharova, T. Y., Rapisarda, V., et al. (2011). p63 regulates Satb1 to control tissue-specific chromatin remodeling during development of the epidermis. J Cell Biol 194, 825–839. doi: 10.1083/jcb.201101148

10. Fisher, M. L., Balinth, S., and Mills, A. A. (2020). p63-related signaling at a glance. J Cell Sci 133. doi: 10.1242/jcs.228015

11. Foffi, E., Violante, A., Pecorari, R., Lena, A. M., Rugolo, F., Melino, G., et al. (2024). BRD4 sustains p63 transcriptional program in keratinocytes. Biology Direct 19, 124. doi: 10.1186/s13062-024-00547-1

12. Ghioni, P., Bolognese, F., Duijf, P. H. G., Van Bokhoven, H., Mantovani, R., and Guerrini, L. (2002). Complex Transcriptional Effects of p63 Isoforms: Identification of Novel Activation and Repression Domains†. Molecular and Cellular Biology 22, 8659–8668. doi: 10.1128/MCB.22.24.8659-8668.2002

13. Guo, Y., Wu, H., Wiesmüller, L., and Chen, M. (2024). Canonical and non-canonical functions of p53 isoforms: potentiating the complexity of tumor development and therapy resistance. Cell Death Dis 15, 1–16. doi: 10.1038/s41419-024-06783-7

14. Heinz, S., Benner, C., Spann, N., Bertolino, E., Lin, Y. C., Laslo, P., et al. (2010). Simple combinations of lineage-determining transcription factors prime cis-regulatory elements required for macrophage and B cell identities. Mol. Cell 38, 576–589. doi: 10.1016/j.molcel.2010.05.004

15. Helton, E. S., Zhang, J., and Chen, X. (2008). The proline-rich domain in p63 is necessary for the transcriptional and apoptosis-inducing activities of TAp63. Oncogene 27, 2843– 2850. doi: 10.1038/sj.onc.1210948

16. Helton, E. S., Zhu, J., and Chen, X. (2006). The unique NH2-terminally deleted (DeltaN) residues, the PXXP motif, and the PPXY motif are required for the transcriptional activity of the DeltaN variant of p63. J Biol Chem 281, 2533–2542. doi: 10.1074/jbc.M507964200

17. Hermeking, H., Lengauer, C., Polyak, K., He, T.-C., Zhang, L., Thiagalingam, S., et al. (1997). 14-3-3σ Is a p53-Regulated Inhibitor of G2/M Progression. Molecular Cell 1, 3–11. doi: 10.1016/S1097-2765(00)80002-7

18. Karsli Uzunbas, G., Ahmed, F., and Sammons, M. A. (2019). Control of p53-dependent transcription and enhancer activity by the p53 family member p63. J. Biol. Chem. doi: 10.1074/jbc.RA119.007965

19. Khan, A., and Mathelier, A. (2017). Intervene: a tool for intersection and visualization of multiple gene or genomic region sets. BMC Bioinformatics 18, 287. doi: 10.1186/s12859-017-1708-7

20. Kim, D., Paggi, J. M., Park, C., Bennett, C., and Salzberg, S. L. (2019). Graph-based genome alignment and genotyping with HISAT2 and HISAT-genotype. Nat Biotechnol 37, 907– 915. doi: 10.1038/s41587-019-0201-4

21. Kouwenhoven, E. N., Oti, M., Niehues, H., van Heeringen, S. J., Schalkwijk, J., Stunnenberg, H. G., et al. (2015). Transcription factor p63 bookmarks and regulates dynamic enhancers during epidermal differentiation. EMBO reports 16, 863–878. doi: 10.15252/embr.201439941

22. LeBoeuf, M., Terrell, A., Trivedi, S., Sinha, S., Epstein, J. A., Olson, E. N., et al. (2010). Hdac1 and Hdac2 act redundantly to control p63 and p53 functions in epidermal progenitor cells. Dev Cell 19, 807–818. doi: 10.1016/j.devcel.2010.10.015

23. Lee, C. T., Cavalcante, R. G., Lee, C., Qin, T., Patil, S., Wang, S., et al. (2020). Poly-Enrich: count-based methods for gene set enrichment testing with genomic regions. NAR Genomics and Bioinformatics 2, lqaa006. doi: 10.1093/nargab/lqaa006

24. Lena, A. M., Rossi, V., Osterburg, S., Smirnov, A., Osterburg, C., Tuppi, M., et al. (2021). The p63 C-terminus is essential for murine oocyte integrity. Nat Commun 12, 383. doi: 10.1038/s41467-020-20669-0

25. Li, D., Kok, C. Y. L., Wang, C., Ray, D., Osterburg, S., Dötsch, V., et al. (2024). Dichotomous transactivation domains contribute to growth inhibitory and promotion functions of TAp73. Proc. Natl. Acad. Sci. U.S.A. 121, e2318591121. doi: 10.1073/pnas.2318591121

26. Li, H., Handsaker, B., Wysoker, A., Fennell, T., Ruan, J., Homer, N., et al. (2009). The Sequence Alignment/Map format and SAMtools. Bioinformatics 25, 2078–2079. doi: 10.1093/bioinformatics/btp352

27. Li, L., Wang, Y., Torkelson, J. L., Shankar, G., Pattison, J. M., Zhen, H. H., et al. (2019). TFAP2C- and p63-Dependent Networks Sequentially Rearrange Chromatin Landscapes to Drive Human Epidermal Lineage Commitment. Cell Stem Cell 24, 271–284.e8. doi: 10.1016/j.stem.2018.12.012

28. Li, Y., Giovannini, S., Wang, T., Fang, J., Li, P., Shao, C., et al. (2023). p63: a crucial player in epithelial stemness regulation. Oncogene 42, 3371–3384. doi: 10.1038/s41388-023-02859-4

29. Li, Y., Zhou, Z., and Chen, C. (2008). WW domain-containing E3 ubiquitin protein ligase 1 targets p63 transcription factor for ubiquitin-mediated proteasomal degradation and regulates apoptosis. Cell Death Differ 15, 1941–1951. doi: 10.1038/cdd.2008.134

30. Lin-Shiao, E., Lan, Y., Welzenbach, J., Alexander, K. A., Zhang, Z., Knapp, M., et al. (2019). p63 establishes epithelial enhancers at critical craniofacial development genes. Science Advances 5, eaaw0946. doi: 10.1126/sciadv.aaw0946

31. Livera, G., Petre-Lazar, B., Guerquin, M.-J., Trautmann, E., Coffigny, H., and Habert, R. (2008). p63 null mutation protects mouse oocytes from radio-induced apoptosis. Reproduction 135, 3–12. doi: 10.1530/REP-07-0054

32. Love, M. I., Huber, W., and Anders, S. (2014). Moderated estimation of fold change and dispersion for RNA-seq data with DESeq2. Genome Biol. 15, 550. doi: 10.1186/s13059-014-0550-8

33. Marshall, C. B., Beeler, J. S., Lehmann, B. D., Gonzalez-Ericsson, P., Sanchez, V., Sanders, M. E., et al. (2021). Tissue-specific expression of p73 and p63 isoforms in human tissues. Cell Death Dis 12, 1–10. doi: 10.1038/s41419-021-04017-8

34. McGrath, J. A., Duijf, P. H. G., Doetsch, V., Irvine, A. D., Waal, R. de, Vanmolkot, K. R. J., et al. (2001). Hay–Wells syndrome is caused by heterozygous missense mutations in the SAM domain of p63. Human Molecular Genetics 10, 221–230. doi: 10.1093/hmg/10.3.221

35. Mills, A. A., Zheng, B., Wang, X. J., Vogel, H., Roop, D. R., and Bradley, A. (1999). p63 is a p53 homologue required for limb and epidermal morphogenesis. Nature 398, 708–13. doi: 10.1038/19531

36. Murray-Zmijewski, F., Lane, D. P., and Bourdon, J.-C. (2006). p53/p63/p73 isoforms: an orchestra of isoforms to harmonise cell differentiation and response to stress. Cell Death Differ 13, 962–972. doi: 10.1038/sj.cdd.4401914

37. Nguyen, B.-C., Lefort, K., Mandinova, A., Antonini, D., Devgan, V., Della Gatta, G., et al. (2006). Cross-regulation between Notch and p63 in keratinocyte commitment to differentiation. Genes Dev 20, 1028–1042. doi: 10.1101/gad.1406006

38. Pecorari, R., Bernassola, F., Melino, G., and Candi, E. (2022). Distinct interactors define the p63 transcriptional signature in epithelial development or cancer. Biochemical Journal 479, 1375–1392. doi: 10.1042/BCJ20210737

39. Pitzius, S., Osterburg, C., Gebel, J., Tascher, G., Schäfer, B., Zhou, H., et al. (2019). TA*p63 and GTAp63 achieve tighter transcriptional regulation in quality control by converting an inhibitory element into an additional transactivation domain. Cell Death Dis 10, 1–14. doi: 10.1038/s41419-019-1936-z

40. Qu, J., Tanis, S. E. J., Smits, J. P. H., Kouwenhoven, E. N., Oti, M., Van Den Bogaard, E. H., et al. (2018). Mutant p63 Affects Epidermal Cell Identity through Rewiring the Enhancer Landscape. Cell Reports 25, 3490–3503.e4. doi: 10.1016/j.celrep.2018.11.039

41. Qu, J., Yi, G., and Zhou, H. (2019). p63 cooperates with CTCF to modulate chromatin architecture in skin keratinocytes. Epigenetics Chromatin 12, 31. doi: 10.1186/s13072-019-0280-y

42. Quinlan, A. R., and Hall, I. M. (2010). BEDTools: a flexible suite of utilities for comparing genomic features. Bioinformatics 26, 841–842. doi: 10.1093/bioinformatics/btq033

43. Rahimov, F., Marazita, M. L., Visel, A., Cooper, M. E., Hitchler, M. J., Rubini, M., et al. (2008). Disruption of an AP-2alpha binding site in an IRF6 enhancer is associated with cleft lip. Nat. Genet. 40, 1341–1347. doi: 10.1038/ng.242

44. Ramírez, F., Ryan, D. P., Grüning, B., Bhardwaj, V., Kilpert, F., Richter, A. S., et al. (2016). deepTools2: a next generation web server for deep-sequencing data analysis. Nucleic Acids Res 44, W160–W165. doi: 10.1093/nar/gkw257

45. Ramsey, M. R., He, L., Forster, N., Ory, B., and Ellisen, L. W. (2011). Physical association of HDAC1 and HDAC2 with p63 mediates transcriptional repression and tumor maintenance in squamous cell carcinoma. Cancer Res 71, 4373–4379. doi: 10.1158/0008-5472.CAN-11-0046

46. Romano, R.-A., Ortt, K., Birkaya, B., Smalley, K., and Sinha, S. (2009). An Active Role of the ΔN Isoform of p63 in Regulating Basal Keratin Genes K5 and K14 and Directing Epidermal Cell Fate. PLoS One 4, e5623. doi: 10.1371/journal.pone.0005623

47. Serber, Z., Lai, H. C., Yang, A., Ou, H. D., Sigal, M. S., Kelly, A. E., et al. (2002). A C-Terminal Inhibitory Domain Controls the Activity of p63 by an Intramolecular Mechanism. Molecular and Cellular Biology 22, 8601–8611. doi: 10.1128/MCB.22.24.8601-8611.2002

48. Sethi, I., Romano, R.-A., Gluck, C., Smalley, K., Vojtesek, B., Buck, M. J., et al. (2015). A global analysis of the complex landscape of isoforms and regulatory networks of p63 in human cells and tissues. BMC Genomics 16, 584. doi: 10.1186/s12864-015-1793-9

49. Su, X., Paris, M., Gi, Y. J., Tsai, K. Y., Cho, M. S., Lin, Y.-L., et al. (2009). TAp63 prevents premature aging by promoting adult stem cell maintenance. Cell Stem Cell 5, 64–75. doi: 10.1016/j.stem.2009.04.003

50. Suh, E.-K., Yang, A., Kettenbach, A., Bamberger, C., Michaelis, A. H., Zhu, Z., et al. (2006). p63 protects the female germ line during meiotic arrest. Nature 444, 624–628. doi: 10.1038/nature05337

51. Tadeu, A. M. B., and Horsley, V. (2013). Notch signaling represses p63 expression in the developing surface ectoderm. Development 140, 3777–3786. doi: 10.1242/dev.093948

52. Thomason, H. A., Zhou, H., Kouwenhoven, E. N., Dotto, G.-P., Restivo, G., Nguyen, B.-C., et al. (2010). Cooperation between the transcription factors p63 and IRF6 is essential to prevent cleft palate in mice. J Clin Invest 120, 1561–1569. doi: 10.1172/JCI40266

53. van Bokhoven, H., Hamel, B. C. J., Bamshad, M., Sangiorgi, E., Gurrieri, F., Duijf, P. H. G., et al. (2001). p63 Gene Mutations in EEC Syndrome, Limb-Mammary Syndrome, and Isolated Split Hand–Split Foot Malformation Suggest a Genotype-Phenotype Correlation. The American Journal of Human Genetics 69, 481–492. doi: 10.1086/323123

54. Welch, R. P., Lee, C., Imbriano, P. M., Patil, S., Weymouth, T. E., Smith, R. A., et al. (2014). ChIP-Enrich: gene set enrichment testing for ChIP-seq data. Nucleic Acids Research 42, e105. doi: 10.1093/nar/gku463

55. Wolff, S., Talos, F., Palacios, G., Beyer, U., Dobbelstein, M., and Moll, U. M. (2009). The α/β carboxy-terminal domains of p63 are required for skin and limb development. New insights from the Brdm2 mouse which is not a complete p63 knockout but expresses p63 γ-like proteins. Cell Death Differ 16, 1108–1117. doi: 10.1038/cdd.2009.25

56. Yalcin-Ozuysal, Ö., Fiche, M., Guitierrez, M., Wagner, K.-U., Raffoul, W., and Brisken, C. (2010). Antagonistic roles of Notch and p63 in controlling mammary epithelial cell fates. Cell Death Differ 17, 1600–1612. doi: 10.1038/cdd.2010.37

57. Yang, A., Kaghad, M., Wang, Y., Gillett, E., Fleming, M. D., Dötsch, V., et al. (1998). p63, a p53 Homolog at 3q27–29, Encodes Multiple Products with Transactivating, Death-Inducing, and Dominant-Negative Activities. Molecular Cell 2, 305–316. doi: 10.1016/S1097-2765(00)80275-0

58. Yang, A., Schweitzer, R., Sun, D., Kaghad, M., Walker, N., Bronson, R. T., et al. (1999). p63 is essential for regenerative proliferation in limb, craniofacial and epithelial development. Nature 398, 714–718. doi: 10.1038/19539

59. Yin, Y., Stephen, C. W., Luciani, M. G., and Fåhraeus, R. (2002). p53 Stability and activity is regulated by Mdm2-mediated induction of alternative p53 translation products. Nat Cell Biol 4, 462–467. doi: 10.1038/ncb801

60. Zhang, Y., Liu, T., Meyer, C. A., Eeckhoute, J., Johnson, D. S., Bernstein, B. E., et al. (2008). Model-based Analysis of ChIP-Seq (MACS). Genome Biology 9, R137. doi: 10.1186/gb-2008-9-9-r137

61. Zhou, Y., Zhou, B., Pache, L., Chang, M., Khodabakhshi, A. H., Tanaseichuk, O., et al. (2019). Metascape provides a biologist-oriented resource for the analysis of systems-level datasets. Nature Communications 10, 1523. doi: 10.1038/s41467-019-09234-6

